# *miR-34/449 miRNAs* regulate choroid plexus ciliogenesis to control cerebrospinal fluid production

**DOI:** 10.64898/2026.08.30.747939

**Authors:** Suifang Mao, Rui Song, Aleksandra Jovanovic, Shibo Jin, Song Pang, Danielle M Jorgens, Michael F. Wendland, Adam Zimmerman, David Lin, Zhenyu Xuan, C Shan Xu, Harald F. Hess, Srigokul Upadhyayula, Lin He

**Affiliations:** Division of Cellular and Developmental Biology, MCB department, University of California at Berkeley, Berkeley, CA 94705, USA; Janelia Research Campus, Howard Hughes Medical Institute, Ashburn, VA 20147, USA; Department of Cellular & Molecular Physiology, Yale School of Medicine, New Haven, CT 06510, USA; Electron Microscope Laboratory, University of California, Berkeley, Berkeley, CA, 94720 USA; Department of Biological Sciences, Center for Systems Biology, University of Texas at Dallas, Richardson, TX 75080, USA; Chan Zuckerberg Biohub, San Francisco, CA, USA; Molecular Biophysics and Integrated Bioimaging Division, Lawrence Berkeley National Laboratory, Berkeley, CA, USA

**Author notes:** These authors contributed equally (Suifang Mao, Rui Song).

## Abstract

A developmental increase in cerebrospinal fluid (CSF) production during development is essential for neuronal growth and ventricular expansion. A key regulator of CSF production is the specialized sensory multicilia of the choroid plexus (ChP), which mediate non-canonical Sonic hedgehog (Shh) signaling to suppress water channel and ion transporter expression, thereby limiting CSF production. ChP multicilia progressively shortens during development, attenuating Shh signaling and promoting CSF production. Here, we identify *miR-34/449* miRNAs as essential regulators of ChP multiciliogenesis. Whereas mutations in canonical ciliogenesis genes elevate CSF production and contribute to hydrocephaly, deletion of *miR-34/449* reduces CSF volume and causes microcephaly. Loss of *miR-34/449* miRNAs causes excessive basal body amplification, defective basal body docking, and failure of developmental multiciliary shortening. Consequently, *miR-34/449*-deficient ChP cilia remain abnormally long and fail to attenuate Shh signaling, resulting in sustained repression of water channel and ion transporter expression and reduced CSF production. Mechanistically, *miR-34/449* miRNAs directly target *Gmnc*, a master transcriptional regulator of multiciliogenesis, to restrain basal body amplification and promote basal body docking. Together, our findings identify *miR-34/449* miRNAs as critical regulators of ChP multiciliogenesis and establish the developmental remodeling of ChP multicilia as a mechanism to couple Shh signaling dynamics to developmental control of CSF production.

## Introduction

The choroid plexus (ChP) is a highly vascularized, secretory neuroepithelium within the brain ventricles, serving as the primary site of cerebrospinal fluid (CSF) production (1–4). The choroid plexus consists of a monolayer of multiciliated choroid plexus epithelial cells (CPECs) that envelop the underlying capillaries and connective tissue, regulating water and ion transport from blood supply into the brain ventricles through a cohort of water channels, ion channels and transporters (1, 2, 5–7). In adults, steady CSF production by choroid plexus is essential for brain homeostasis, supporting mechanical protection, nutrient delivery, and waste clearance (8, 9). Yet during embryonic and postnatal development, a rapid increase in CSF production generates ventricular hydrostatic pressure to facilitate the expansion of brain ventricles (10–15). Hence, impaired CSF production due to choroid plexuses defects give rise to brain developmental and physiological phenotype, such as hydrocephalus, neurodegeneration, and neuroinflammation (16–20).

The multi sensory cilia on CPECs are key cellular structures that govern the regulation of CSF production (18, 21, 22). Cilia are microtubule-based cellular protrusions that integrate cellular signaling, motility, and environmental sensing across a variety of cell types (23, 24). Choroid plexus multicilia are a unique class of sensory multicilia that combine ultrastructural features of both primary and motile cilia while also possessing distinctive structural characteristics (18, 25). Each CPEC contains roughly 18 “9+0” sensory cilia per cell. Choroid plexus cilia mediate a non-canonical Shh signaling to downregulate cAMP, and subsequently repress the expression of the water channel Aqp1 and the ion transporter Atp1a2 to dampen CSF production (18). Fewer and shorter choroid plexus cilia caused by deficiency of core ciliogenesis regulators, such as *Foxj1* and *Ift88*, attenuate Shh signaling, derepress water channel and ion transporter expression, increase CSF production, and contribute to hydrocephalus (18). During embryonic and postnatal development when the brain ventricles rapidly expand in size, the choroid plexus ciliary length is dynamically decreased. This dynamic ciliary length decrease attenuates Shh signaling, elevates water channel and ion transporter expression, and ultimately, promotes a developmental increase in CSF production (18). This precise and dynamic regulation of CSF production is critical to maintain appropriate intracranial pressure and ventricular architecture as the brain ventricles expand (12, 13, 15).

Here, we identify *miR-34/449* miRNAs as key regulators for choroid plexus ciliogenesis. miRNAs are small regulatory non-coding RNAs that regulate a large cohort of mRNA targets posttranscriptionally (26–29). The *miR-34/449* family comprises six evolutionarily conserved mature miRNAs (*miR-34a*, *miR-34b*, *miR-34c*, *miR-449a*, *miR-449b* and *miR-449c*), encoded by three genomic loci (*miR-34a, miR-34b/c* and *miR-449a/449b/449c*) (30–32). *miR-34/449* miRNAs are highly enriched in ciliated cells, including airway multiciliated cells (MCCs), which contain hundreds of apical motile cilia per cell; sperms, which contain a single motile flagellum; and CPECs, which contain ∼18 centrally positioned sensory multicilia per cell (33, 34). *miR-34/449*-deficient mice exhibit classic motile cilia defects in the airway and reproductive tract, including impaired mucociliary clearance and male infertility, similar to those observed in mice lacking canonical ciliogenesis genes such as *Foxj1* and *Ift88*. (33–37). However, in contrast to *FoxJ1^−/−^* or *Ift88^−/−^* mutants that often develop hydrocephalus due to impaired CSF flow and increased CSF production (18, 38, 39), *miR-34/449*–deficient mice exhibit microcephaly associated with reduced CSF production by choroid plexus. *miR-34/449* deficient CPECs display increased cilia and basal body numbers, aberrant basal body docking and elongated ciliary axonemes. These ciliogenesis defects cause the failure to trigger developmental decrease in ciliary length, resulting in elevated basal Sonic hedgehog (Shh) signaling, a sustained repression of specific water/ion channels and transporters, and decreased CSF production and microcephaly.

Mechanistically, *miR-34/449* miRNAs act, at least in part, by dampening the expression of Gmnc, a central transcription factor for multiciliogenesis. The precise regulation of Gmnc level in CPECs plays an important role in basal body amplification and ChP cilia dynamics. The defective basal body docking in *miR-34/449* deficient CPECs disrupts ChP multicilia shortening. Hence, *miR-34/449* miRNAs are key regulators for choroid plexus multiciliogenesis, linking ciliary dynamics to the developmental modulation of Shh signaling and CSF production.

## Results

### *miR-34/449* miRNAs are highly enriched in multiciliated CPECs

*miR-34/449* miRNAs are key regulators of motile ciliogenesis, with highly enriched expression in airway epithelia, testis, and ependymal epithelia (33). Similar to mutants lacking canonical motile ciliogenesis genes, *miR-34/449*-deficient mice exhibit primary ciliary dyskinesia-like phenotypes, including postnatal mortality, infertility, and defective mucociliary clearance (33–35). Surprisingly, *miR-34/449*-deficient mice do not develop hydrocephalus (33), a common phenotype of canonical motile ciliogenesis mutants that results from impaired ependymal cilia function and disrupted CSF circulation. (40–43). This unexpected phenotypic divergence suggests that *miR-34/449* regulates multiciliogenesis through mechanisms distinct from those controlled by canonical ciliogenesis genes.

We quantified individual *miR-34/449* miRNAs across different brain regions by real-time PCR, and found the highest expression level in the choroid plexus during late embryonic and postnatal development, markedly exceeding levels in ventricular regions enriched for ependymal motile cilia (Fig. 1A). *In situ* hybridization analyses on *miR-34a, miR-34c* and *miR-449a* demonstrated their enrichment in CPECs, a ciliated cell type with sensory multicilia (Fig. 1B). *miR-34/449* enrichment in ciliated cells with sensory multicilia (such as CPECs) and motile multicilia (such as airway MCC) implicates their functional importance across multiple multiciliated cell types.

**Figure 1.**
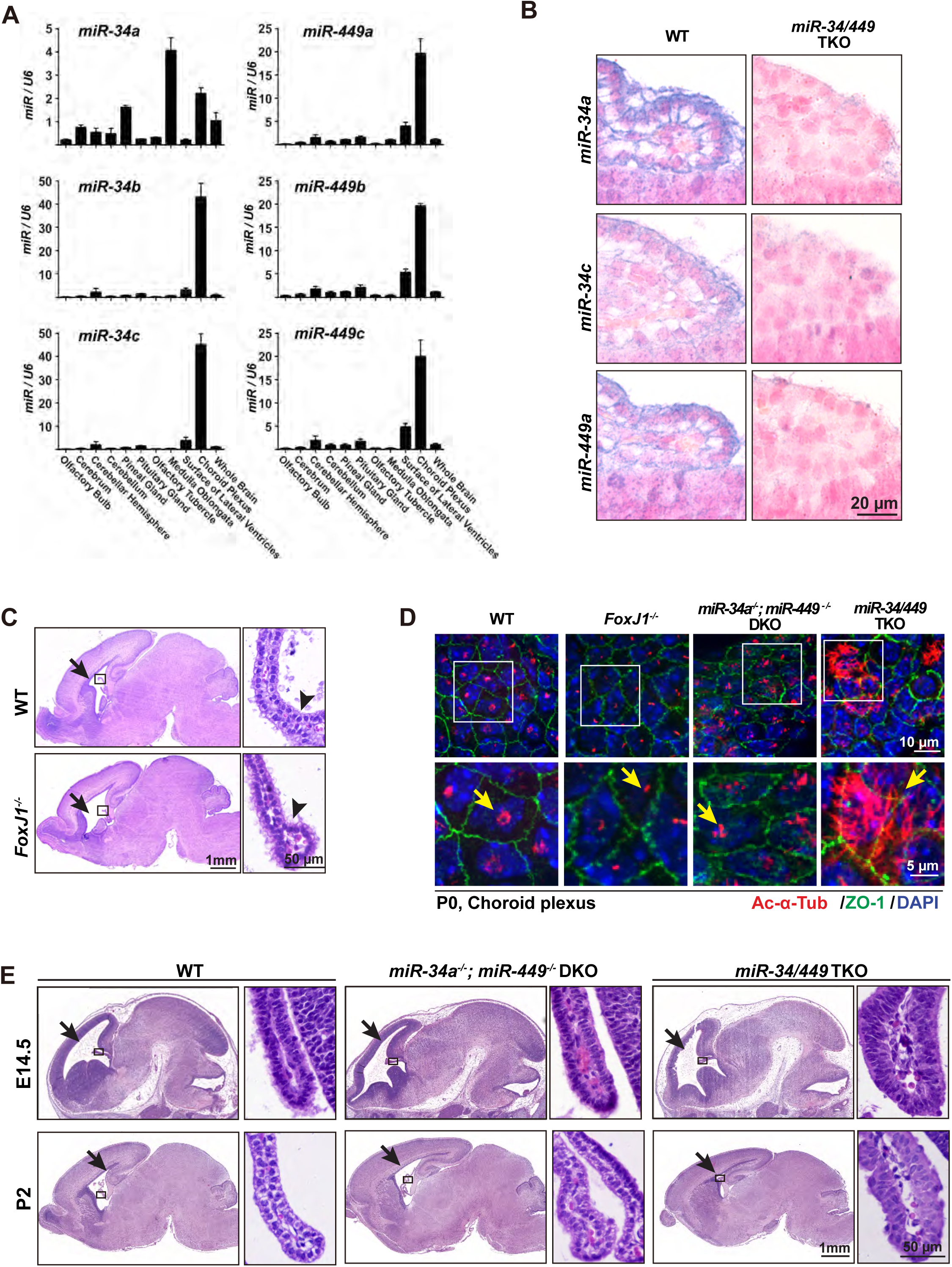
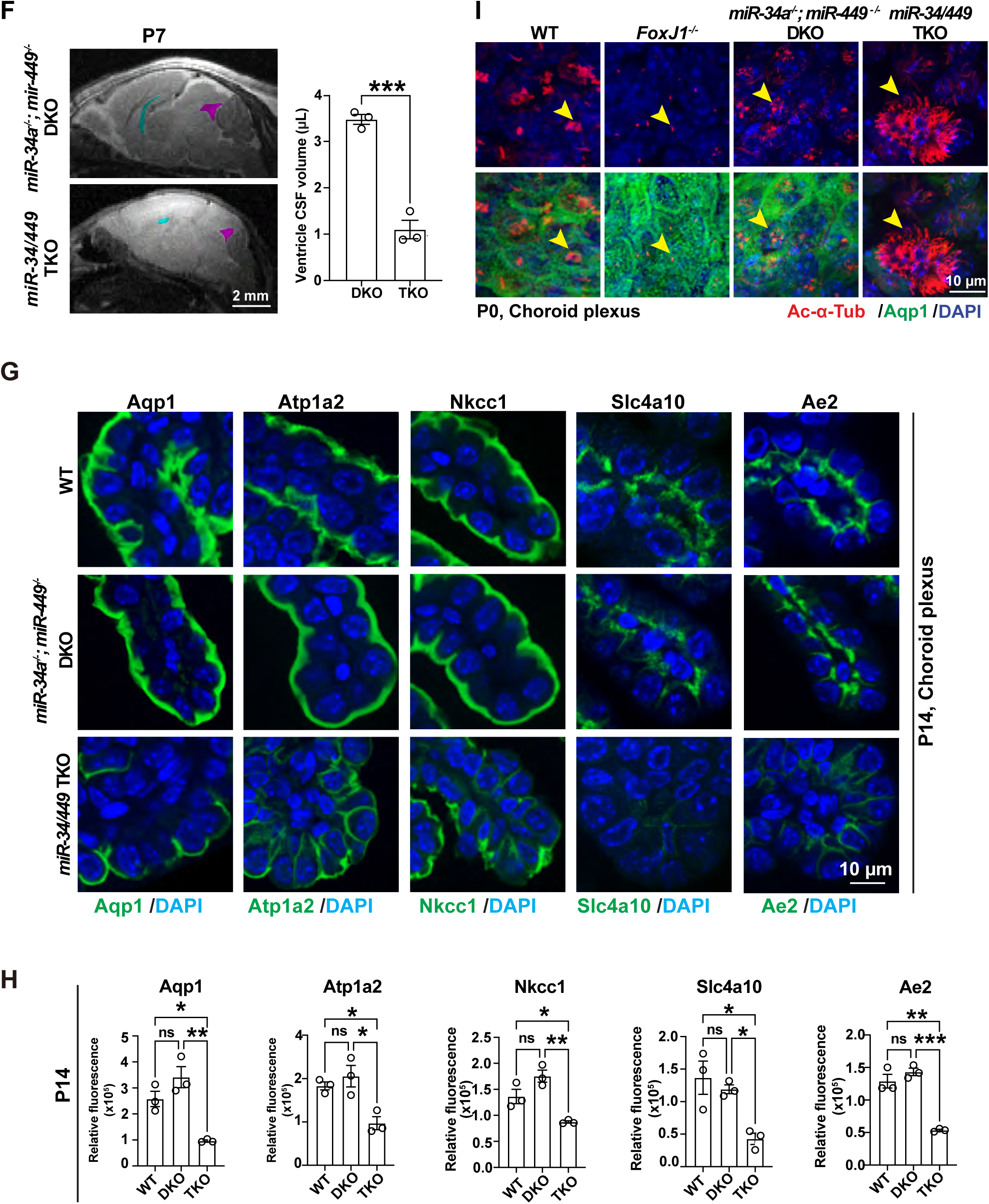
Deletion of *miR-34/449* miRNAs in mice decreases CSF production and cause microcephaly. **A, B.** *miR-34/449 miRNAs* are enriched in choroid plexus epithelial cells (CPECs). **A.** The expression of *miR-34a*, *miR-34b, miR-34c*, *miR-449a, miR-449b* and *miR-449c* miRNAs were measured in multiple dissected brain regions using real time PCR analysis. **B**. *In situ* hybridization analyses revealed the enrichment of *miR-34a, miR-34c* and *miR-449a* miRNAs in wide type CPECs, but not *miR-34/449* TKO choroid plexus. **C.** *FoxJ1* deficiency in P3 mice gives rise to enlarged lateral ventricles (arrows) and a more compacted morphology for CPECs (arrowheads) in histology analysis**. D.** *miR-34/449* eficiency increases cilia number and ciliary length in P0 CPECs and *FoxJ1* deficiency has the opposite cilia phenotype. Representative immunostaining images were shown for P0 littermate controlled wildtype (WT) and *FoxJ1^−/−^* ChP (n=3) and P0 littermate controlled *miR34a*^−/−^; *miR449*^−/−^ DKO and *miR-34/449* TKO ChP (n=3). Ac-α-Tub (ciliary axoneme), ZO-1 (tight junction) and DAPI. **E.** *miR-34/449* deficiency decreases the size of lateral ventricles (arrows) in mice (left) and alters the cell morphology of CPECs (right). Littermate controlled *miR34a*^−/−^; *miR449*^−/−^ DKO and *miR-34/449* TKO mice, along with aged matched WT mice, were compared at E14.5 (n=3), and P2 (n=3). **F.** MRI reveals decreased CSF volume in the lateral ventricles and the 4th ventricles in P7 *miR-34/449* TKO mice. Sagittal sections of MRI scan (left) and quantitative measurement of CSF volume (right) were shown for P7 littermate-controlled *miR34a*^−/−^; *miR449*^−/−^ DKO and *miR-34/449* TKO mice (n=3). Cyan, lateral ventricles; magenta, the fourth ventricles. Total ventricle CSF volume, n=3, P7 DKO vs. P7 TKO, \*\*\**P* = 0.0005, t = 10.57, df = 4; Error bars, sem; unpaired two-tailed Student’s t-test. **G, H.** *miR-34/449* deficiency decreases the expression of water channels and ion transporters in P14 choroid plexus. Representative immunostaining images (**G**) and quantitation of relative fluorescence (**H**) were shown for Aqp1, Atp1a2, Nkcc1, Slc4A10 and Ae2 in P14 wildtype, *miR34a*^−/−^; *miR449*^−/−^ DKO and *miR-34/449* TKO choroid plexus paraffin sections (n=3 for each genotype). In **H**, each dot is the average intensity from 3-6 images in one choroid plexus sample. The statistical tests used in each graph are listed in Supplementary Table S4, nested one-way ANOVA and Sidak multiple comparisons tests. **I.** Immunostaining on whole-mount P0 ChP tissue showed a decrease in apical Aqp1 expression in *miR-34/449* TKO CPECs, but an increase in *FoxJ1^−/−^* CPECs. Yellow arrowheads, CPECs. Three pairs of littermate-controlled *miR34a*^−/−^; *miR449*^−/−^ DKO and *miR-34/449* TKO mice, along with aged matched WT mice, were examined.

### *miR-34/449* deficient mice exhibit microcephaly with impaired CSF production

The choroid plexus sensory multicilia play an essential role in regulating the extent of CSF production by mediating Sonic hedgehog (Shh) signaling (*Mao et al*., 2025). The key pathways that regulate the ciliogenesis of motile multicilia also govern that of choroid plexus sensory multicilia (18, 22, 44). For example, *FoxJ1* deletion gives rise to fewer and shorter cilia in CPECs, causing impaired Shh signaling and increased CSF production, which contributes to a hydrocephalus phenotype (Fig. 1C). In comparison, while *miR-34/449* triple knockout (TKO) mice also impair choroid plexus ciliogenesis, they exhibit an opposite ciliary phenotype, characterized by a greater number of cilia per cell with an increased ciliary length, and ultimately, microcephaly (Fig. 1D, 1E).

*miR-34/449* TKO mice did not display any obvious brain phenotype at E14.5, when choroid plexus forms a distinct, highly vascularized epithelial structure (Fig. 1E). Yet starting from P2 to P14, when choroid plexus functionally matures and brain ventricles rapidly expand, *miR-34/449* TKO mice exhibit microcephaly (Fig.1E, Sup Fig. S1A), with a reduction in brain size, brain ventricular size and CSF volume (Fig. 1E, 1F). *miR-34/449* deficient CPECs also exhibit an enlarged cell size with a compact morphology (Fig.1E, Sup Fig. S1B, S1C).

### *miR-34/449* deficiency impairs water channel and ion transporter expression

A critical step in CSF production is the directional transport of water and ions from the choroidal blood supply into the brain ventricles, regulated by the polarized expression of specific water and ion channels/transporters in CEPCs (8). While the expression of water channels and ion transporters remained normal at E14.5 in *miR-34/449* TKO ChP (Sup Fig. S1D), we observed a marked decrease by P14, affecting the apically localized Aqp1 (a water channel), Atp1a2 (a subunit of Na^+^/K^+^-ATPase) and Nkcc1 (a Na^+^-K^+^-2Cl^−^ cotransporter), as well as the basolaterally localized Slc4a10 (a Na^+^-dependent HCO3^−^-transporter) and Ae2 (a Cl^−^-HCO3^−^-exchanger) (Fig. 1G, 1H). This reduction occurs at both the mRNA and the protein level in *miR-34/449* TKO ChP (Fig. 1G, 1H, S1E), with the phenotype being most pronounced at P14, coinciding with the severe microcephaly phenotype and reduced CSF production.

Choroid plexus sensory multicilia plays an essential role to regulate the dynamic expression of water channels and ion transporter expression during embryonic and postnatal development. Given the specific enrichment of *miR-34/449* miRNAs in CPECs and their reported effects in regulating motile multiciliogenesis, we examined *miR-34/449* TKO choroid plexus cilia and observed an increase in sensory multicilia number and ciliary axoneme length at P0 (Fig. 1D), which precisely correlated with a decrease in Aqp1 expression (Fig. 1I). Interestingly, the *miR-34/449* TKO choroid plexus phenotype contrasts with that caused by *FoxJ1* deletion, which is characterized by fewer and shorter cilia, elevated Aqp1 expression and hydrocephaly (Fig. 1D, 1I). Hence, the ciliary defects in choroid plexus likely give rise to dysregulated expression of water channels and ion transporters, and ultimately microcephaly, in *miR-34/449* TKO mice.

### *miR-34/449* miRNAs regulate multiciliogenesis in choroid plexus epithelial cells

In contrast to airway MCCs, whose motile multicilia elongate their axonemes during maturation (Fig. 2A), choroid plexus epithelial cells (CPECs) initially possess long, scattered multicilia at E14.5 that progressively mature into short, densely clustered multicilia by P14 (Fig. 2B). E14.5 to P14 is a critical developmental window marked by a rapid increase in CSF production and a substantial expansion of the brain ventricles (18, 25, 45). Focused ion beam–scanning electron microscopy (FIB-SEM) further confirmed the developmental regulation of choroid plexus ciliary length and basal body organization (Fig. 2C). In E14.5,the long and scattered choroid plexus cilia corresponds to the less organized basal bodies. By P0 to P14, basal bodies become centrally clustered through multiple SDA–SDA (subdistal appendage) or SDA–basal body contacts (Fig. 2C, Sup Fig. S2A), assembling into either a single compact, short cilia cluster with tightly bundled axonemes, or multiple closely grouped short cilia subclusters with axonemes grouped into several distinct bundles (Fig. 2C, Sup Fig. S2A).

**Figure 2.**
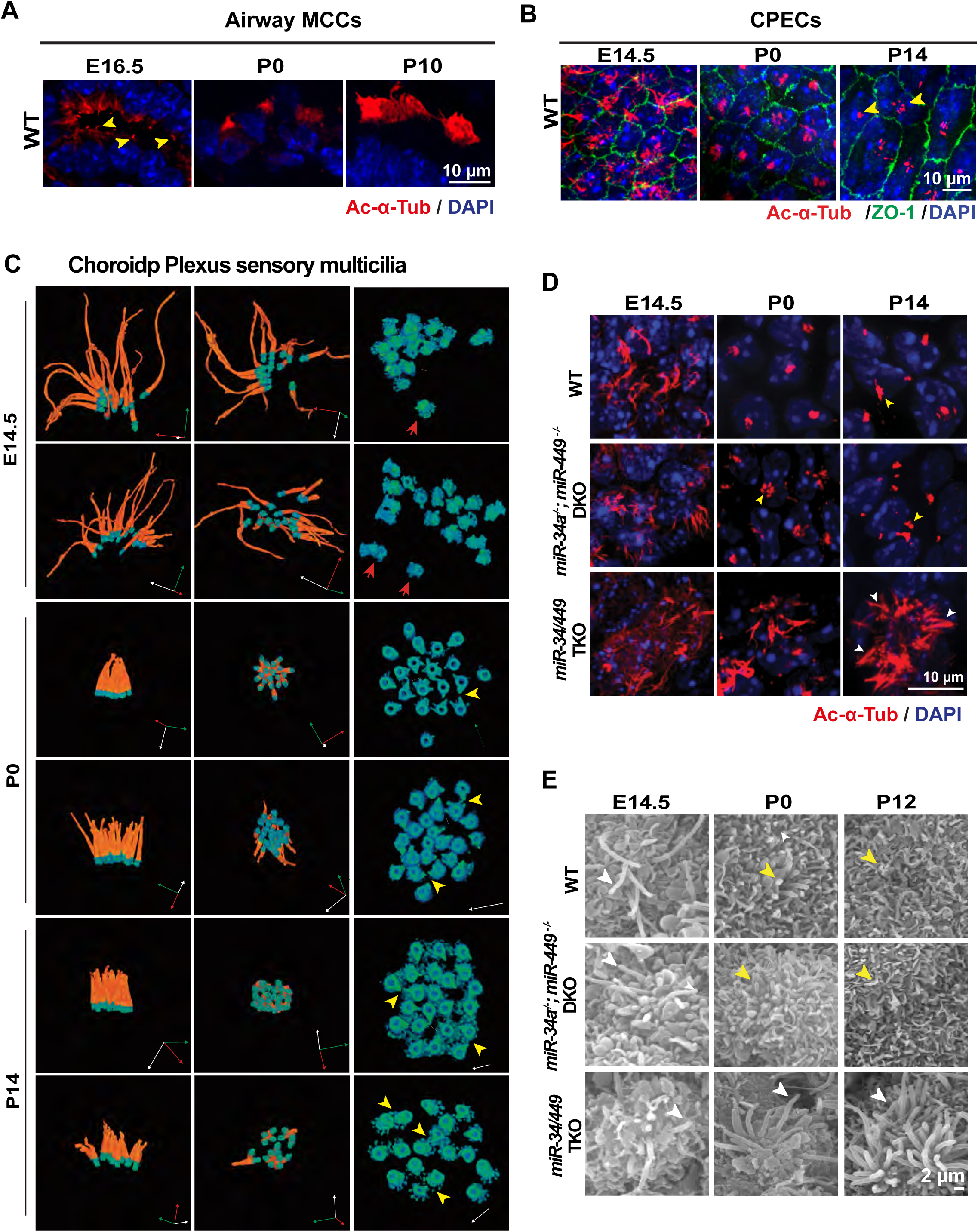
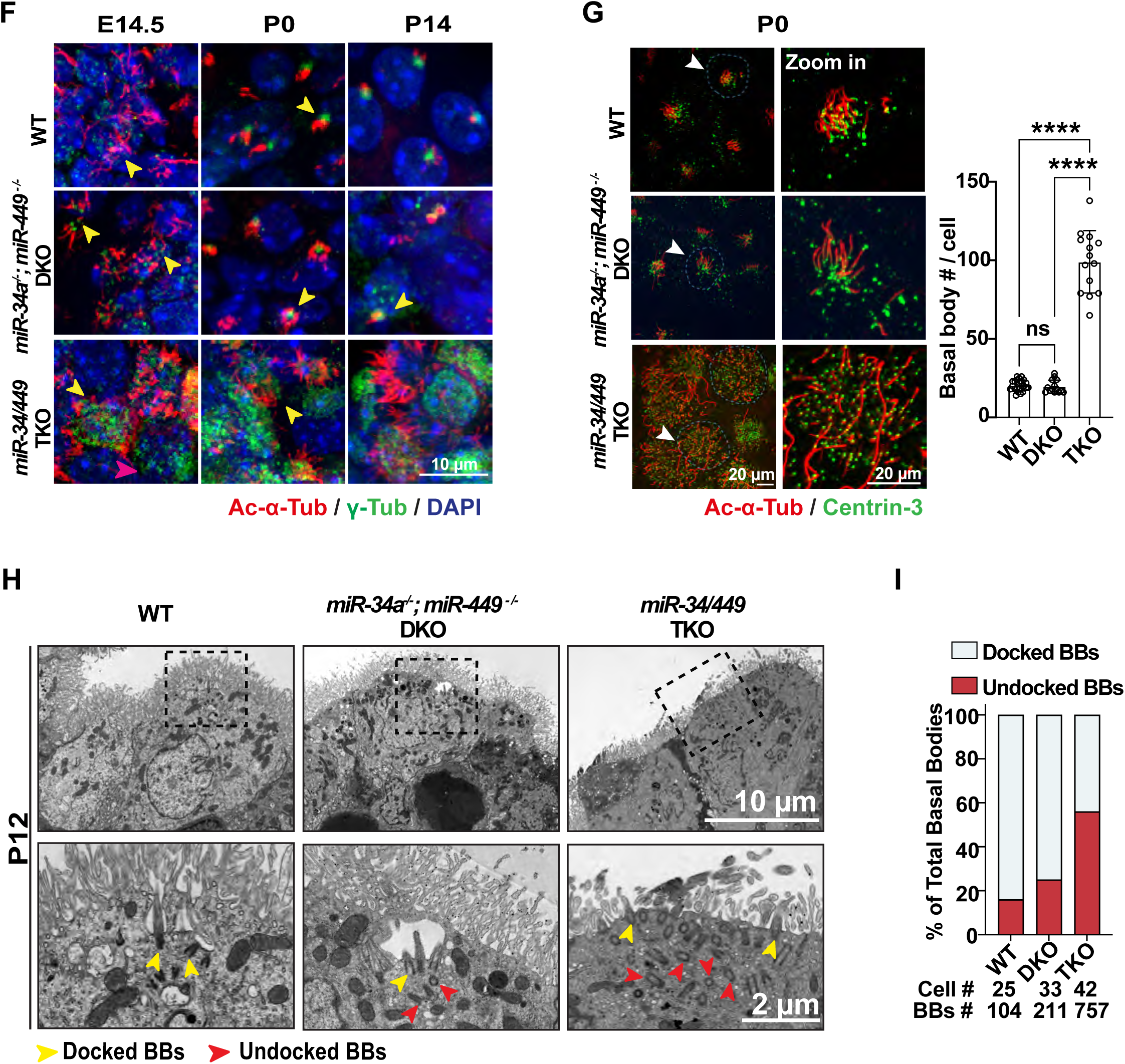
*miR-34/449* deficiency promotes aberrant multiciliogenesis in choroid plexus. **A.** Airway multiciliated cells elongate ciliary axonemes (yellow arrowheads) during maturation, as shown by immunostaining of Ac-α-Tub (ciliary axonemes) and DAPI in lung sections. **B, C.** Choroid plexus multicilia develops from cilia with long and scattered axonemes at E14.5 to those with short, clustered ciliary axonemes at P14. **B.** Representative images of whole-mount tissue immunostaining of Ac-α-Tub (ciliary axonemes), ZO1 (apical tight junction) and DAPI were shown. **C.** Pseudo-color images are shown from FIB-SEM 3D reconstructions of wildtype CPECs in E14.5, P0, and P14. E14.5 choroid plexus cilia are initially characterized by long axonemes (orange) and dispersed basal bodies (blue green) at E14.5. Cilia start to shorten in length and basal bodies start to cluster at P0. By P14, choroid plexus cilia either form a single cluster of short cilia with their basal bodies closely aggregated (top) or form several small clusters with less aggregated basal bodies (bottom). 3D axes, X (red), Y (green), and Z (white). **D-H.** *miR-34/449* deficiency significantly increases cilia length and cilia number in CPECs. **D.** Littermate-controlled E14.5, P0 and P14 *miR-34a^−/−^; miR-449^−/−^* DKO and *mir-34/449* TKO mice, along with age matched WT mice, were subjected to immunostaining for Ac-α-Tub (ciliary axonemes) and DAPI in whole mount tissues. **E.** SEM analysis reveals that cilia in *miR-34/449* TKO choroid plexus are longer (white arrowheads) compared to those in wildtype and *miR-34a^−/−^; miR-449^−/−^* DKO choroid plexus (yellow arrowhead). **F, G.** *miR-34/449* TKO CPECs have an increased number of basal bodies with a scattered apical distribution. Littermate-controlled *miR-34a^−/−^; miR-449^−/−^* DKO and *miR-34/449* TKO choroid plexus tissue (n=3), along with age matched WT tissue, were co-stained for γ-Tubulin (basal body) and Ac-α-Tubulin (ciliary axoneme) for whole-mount tissue immunostaining (**F**), and were co-stained for Ac-α-Tubulin (ciliary axoneme) and Centrin3 (basal body) for expansion microscopy (**G**). Yellow arrowheads in **F** and white arrowheads in **G**, individual CPECs. **G.** Quantitation of basal body number, WT vs. DKO, n.s., *P* = 0.9958, t = 0.2042, df = 48; WT vs. TKO, **** *P* <0.0001, t = 21.47, df = 48; DKO vs. TKO, **** *P* <0.0001, t = 19.44, df = 48; one-way ANOVA and Sidak multiple comparisons tests. Error bars, s.d. **H.** TEM analysis revealed an increased number of undocked basal bodies (BBs) in *miR-34/449* TKO choroid plexus at P12. Red arrowheads, undocked BBs; yellow arrowheads, docked BBs. **I.** Quantitation of docked and undocked BBs in WT, *miR-34a^−/−^; miR-449^−/−^* DKO and *miR-34/449* CPECs.

Compared to wildtype and *miR-34a^−/−^; miR-449^−/−^*DKO controls, the *miR-34/449* TKO choroid plexus exhibits a greater cilia number per cell, impaired ciliary clustering, and elongated ciliary axonemes, as revealed by immunostaining and scanning electron microscopy (SEM) (Fig. 2D, 2E, Sup Fig. S2B). Notably, the earliest detectable ciliary defect in *miR-34/449* TKO choroid plexus is an increase in basal body number, which emerges at E14.5 (Fig. 2F), preceding the onset of all other *miR-34/449* dependent phenotypes. During subsequent postnatal development, *miR-34/449* TKO CPECs exhibit defective cilia clustering, and increased ciliary axoneme number and length (Fig. 2G-2I Sup Fig. S2B). While the majority of the basal bodies in wildtype and *miR-34a^−/−^; miR-449^−/−^* DKO CPECs dock successfully at the apical membrane, only 43.6% basal bodies in *miR-34/449* TKO CPECs are properly docked (Fig. 2H, 2I), and mislocalized basal bodies frequently fail to extend cilia (Fig. 2H, Sup Fig. S2C). Interestingly, the increased basal body amplification outweighs the basal body docking defects, ultimately resulting in an increased number of cilia per cell in *miR-34/449* TKO CPECs (Fig. 2E, 2F, Sup Fig. S2B). Altogether, the ciliary defects in the *miR-34/449-*deficient choroid plexus precede the emergence of all other phenotypes and are likely the primary defects caused by *miR-34/449* deletion.

Both *miR-34/449* TKO and control CPECs exhibit long, scattered cilia in E14.5. While choroid plexus multicilia in wildtype and *miR-34a^−/−^; miR-449^−/−^* DKO mice mature into densely clustered short cilia by P14, *miR-34/449* TKO choroid plexus cilia retain a scattered organization and fail to decrease their ciliary length (Fig. 2D, 2F). This dynamic change in choroid plexus ciliary length is important to attenuate Shh signaling during development and derepress Aqp1 and Atp1a2 expression, thereby increasing CSF production (18). Our findings suggest that *miR-34/449-*mediated regulation of basal body amplification and docking is required for this dynamic ciliary remodeling process to decrease ciliary length in a critical developmental window. Subsequently, Failure of developmental ciliary shortening may contribute to altered signaling pathways and reduced CSF production and, consequently, microcephaly in *miR-34/449* TKO mice.

### Shh signaling is aberrantly elevated in *miR-34/449* deficient CEPCs

Choroid plexus sensory multicilia mediate canonical and non-canonical Shh signaling to regulate choroid plexus development and suppress CSF production (18, 46–48). In particular, choroid plexus multicilia mediate non-canonical Shh signaling that activates Smo to repress cyclic AMP (cAMP) production, thereby suppressing specific water channel and ion transporter expression and downregulating CSF production (18). Hence, impaired choroid plexus ciliogenesis in *miR-34/449* deficient mice likely results in aberrant Shh signaling to reduce CSF production.

We employed an explant culture system to compare the responses of wildtype, *miR-34a^−/−^; miR-449^−/−^* DKO, and *miR-34/449* TKO choroid plexus to exogenous Shh treatment *in vitro*, using *Gli1* expression and ciliary Smo translocation as readouts of the Shh pathway activity. Exogenous Shh robustly induced *Gli1* expression via a Smo-dependent mechanism in P0 wildtype and *miR-34a^−/−^; miR-449^−/−^*DKO choroid plexus explants (Fig. 3A, Sup Fig. S3A, S3B). Shh induced *Gli1* induction depends on Shh-stimulated ciliary Smo translocation (Fig. 3B), as treatment with the Smo inhibitor KAAD-cyclopamine abolishes this induction (Fig. 3A). In contrast, *miR-34/449* TKO choroid plexus explants exhibited elevated *Gli1* levels prior to exogenous Shh treatment (Fig. 3C). This *Gli1* induction coincides with pre-existing Smo enrichment within ciliary axonemes (Fig. 3B, Sup Fig.S3C), and can be repressed by Smo inhibition (Fig. 3A). The basal Shh signaling intensity in *miR-34/449* TKO explants, measured by either *Gli1* expression or ciliary Smo translocation, is comparable to that of exogenous Shh-treated control choroid plexus explants (Fig. 3A, Sup Fig.S3A-S3C). Interestingly, exogenous Shh treatment failed to further induce *Gli1* expression or trigger additional Smo translocation in *miR-34/449* TKO choroid plexus explants (Fig. 3A, 3B, Sup Fig. S3C). Yet pretreatment of *miR-34/449* TKO explant with KAAD-cyclopamine to eliminate ciliary Smo localization restores their responsiveness to exogenous Shh stimulation (Fig. 3D). These findings suggest that Shh signaling is aberrantly activated in *miR-34/449* TKO choroid plexes explants, rendering them unresponsive to exogenous Shh stimulation unless their elevated basal Shh signaling is first dampened. Hence, *miR-34/449* deficiency in choroid plexus likely aberrantly elevates basal Shh signaling in explant culture to a level comparable to maximal exogenous ligand–induced Shh activation in wildtype choroid plexus.

**Figure 3.**
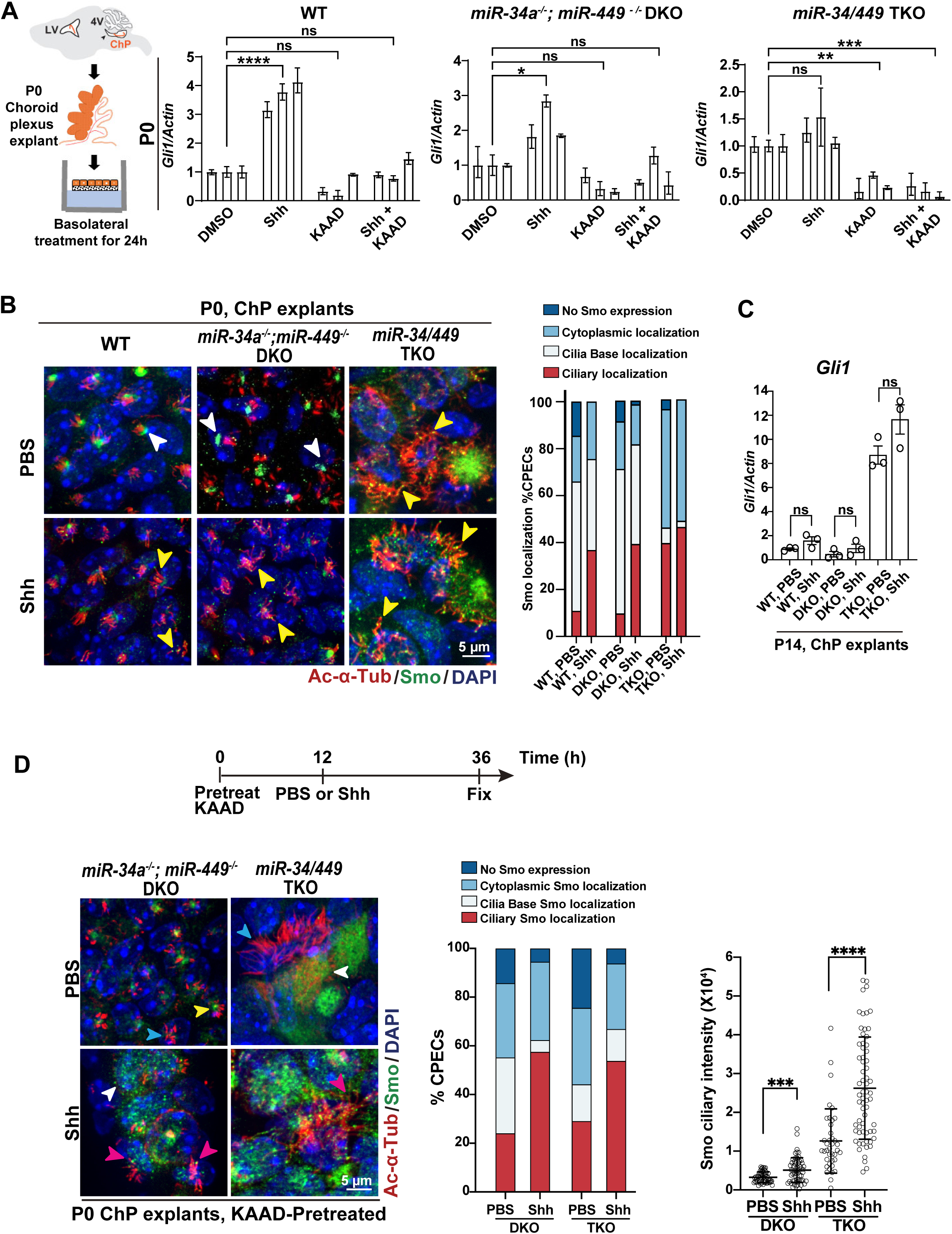
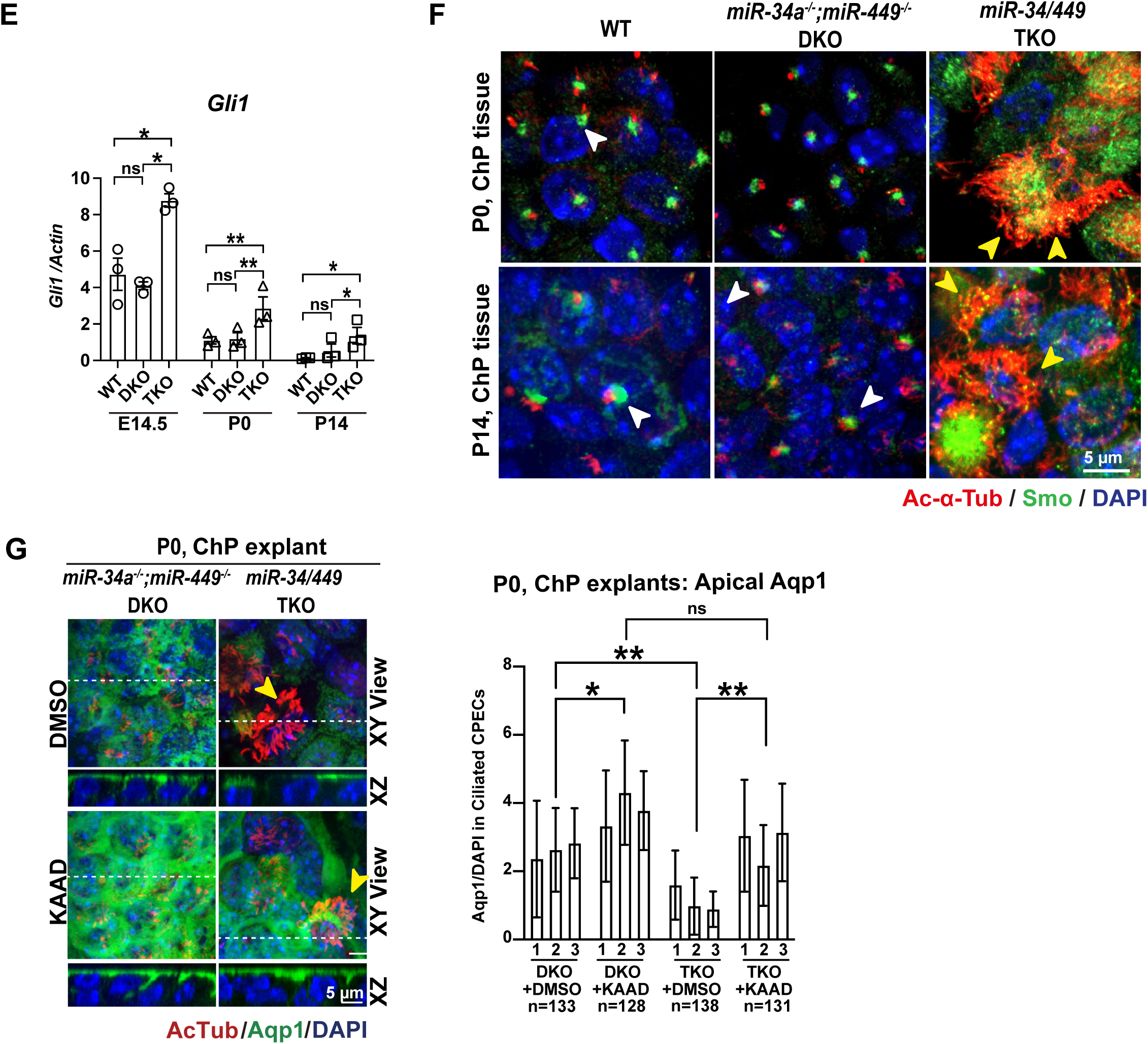
Elongated *miR-34/449* TKO ChP cilia aberrantly elevate the Shh signaling. **A-D**. *miR-34/449* deficiency causes aberrant activation of Shh signaling in the choroid plexus. **A**. In a choroid plexus explant culture system (left), Shh treatment significantly induced *Gli1* expression in a Smo-dependent mechanism in wildtype or *miR-34a^−/−^; miR-449^−/−^* DKO explants. *miR-34/449* choroid plexus failed to respond to exogenous Shh, yet the basal Gli expression could be repressed by Smo inhibition with KAAD treatment. Three pairs of littermate-controlled P0 *miR-34a^−/−^; miR-449^−/−^* DKO and *miR-34/449* TKO choroid plexus, along with age-matched WT explants, were examined, and *Gli1* expression was measured by real time PCR. For WT explants, DMSO vs. Shh, **** *P* < 0.0001, t = 10.52, df = 8; DMSO vs. KAAD, n.s., *P* = 0.0806, t = 2.687, df = 8; DMSO vs. Shh + KAAD, n.s., *P* = 0.9981, t = 0.1618, df = 8. For *miR-34a^−/−^; miR-449^−/−^* DKO explants, DMSO vs. Shh, * *P* = 0.0182, t = 3.693, df = 8; DMSO vs. KAAD, n.s., *P* = 0.2766, t = 1.845, df = 8; DMSO vs. Shh + KAAD, n.s., *P* = 0.8200, t = 0.8211, df = 8. For *miR-34/449* TKO samples, DMSO vs. Shh, n.s., *P* = 0.1568, t = 2.242, df = 8; DMSO vs. KAAD, ** *P* = 0.0013, t = 5.744, df = 8; DMSO vs. Shh + KAAD, *** *P* = 0.0004, t = 6.727, df = 8. One-way ANOVA and Sidak multiple comparisons tests were performed here; error bars, s.d. **B, C.** *miR-34/449* deficient choroid plexus exhibits elevated basal Shh signaling in explant culture, but is unresponsive to exogenous Shh stimulation. Upon Shh treatment, WT and *miR-34a^−/−^; miR-449^−/−^* DKO choroid plexus explants displayed increased ciliary Smo accumulation at P0 (**B**) and elevated *Gli1* expression at P14 (**C**), as assessed by immunostaining and real time PCR, respectively. In contrast, *miR-34/449* TKO choroid plexus exhibited elevated basal ciliary Smo localization and *Gli1* expression prior to Shh treatment, and exogenous Shh failed to further increase Shh pathway activity. In **C**, P14 WT: PBS vs. Shh, n.s., *P* = 0.11, t = 2.047, df = 4; P14 *miR-34a^−/−^; miR-449^−/−^*DKO: PBS vs. Shh, n.s., *P* = 0.302, t = 1.184, df = 4; P14 *miR-34/449* TKO: PBS vs. Shh, n.s., *P* = 0.1066, t = 2.075, df = 4; n =3; unpaired two-tailed Student’s t-test. Error bars, sem. **D**. *miR-34/449* TKO choroid plexus cilia restored their response to exogenous Shh stimulation after a pre-treatment with Smo inhibitor, cyclopamine KAAD. Representative images (left), quantitation of Smo staining patterns (middle), and quantitation of ciliary Smo staining intensity per cell (right) were shown for KAAD-pretreated, P0 *miR-34a^−/−^; miR-449^−/−^* DKO and *miR-34/449* TKO choroid plexus explants (n=3). *miR-34a^−/−^; miR-449^−/−^* DKO explants: PBS (n=54) vs. Shh (n=61), *** *P* =0.0001, t = 3.961, df = 113; *miR-34/449* TKO choroid plexus explants: PBS (n=39) vs. Shh (n=65), **** *P* < 0.0001, t = 5.8, df = 102; unpaired two-tailed Student’s t-test. Error bars, sd. Magenta arrowheads, Smo in ciliary axonemes; yellow arrowheads, Smo at ciliary base; white arrowheads, dispersed Smo in the cytosol; blue arrowheads, a CPEC without Smo staining. **E, F.** *miR-34/449* deficient choroid plexus exhibits elevated Shh signaling *in vivo*. **E.** Real-time PCR analyses of *Gli1* in WT, *miR-34a^−/−^; miR-449^−/−^* DKO, and *miR-34/449* TKO choroid plexus indicated that *Gli1* expression peaks at E14.5 and gradually decreased at P0 and further decreased at P14, and that *Gli1* expression was consistently higher in *miR-34/449* TKO mice at all developmental stages. n=3; error bars, sem, the statistical tests used are listed in Supplementary Table S4, nested one-way ANOVA and Sidak multiple comparisons tests. **F.** Immunostaining for Smo revealed increased ciliary accumulation in the choroid plexus of P0 and P14 *miR-34/449* TKO mice compared with age-matched WT and *miR-34a^−/−^; miR-449^−/−^* DKO littermate controls. **G.** Smo inhibition by KAAD treatment restored the apical Aqp1 expression in *miR-34/449* TKO CPECs. P0 *miR-34a^−/−^; miR-449^−/−^* DKO and *miR-34/449* TKO choroid plexus explants were treated with DMSO or KAAD, before being subjected to immunostaining (left) and quantitation (right) for Aqp1. *miR-34a^−/−^; miR-449^−/−^* DKO DMSO vs. *miR-34/449* TKO DMSO, \*\**P* = 0.0095, t = 4.154, df = 8; *miR-34a^−/−^; miR-449^−/−^* DKO DMSO vs. *miR-34a^−/−^; miR-449^−/−^* DKO KAAD, \**P* = 0.019, t = 4.154, df = 8; *miR-34/449* TKO DMSO vs. *miR-34/449* TKO KAAD, ** *P* = 0.0099, t = 4.648, df = 8; *miR-34a^−/−^; miR-449^−/−^* DKO KAAD vs. *miR-34/449* TKO KAAD, n.s., *P* = 0.1041, t = 2.959, df = 8; nested unpaired One-way ANOVA and Sidak multiple comparison. Error bars, sd.

During embryonic and postnatal development, choroid plexus cilia undergo a reduction in axonemal length and become less responsive to Shh signaling (Sup Fig. S3A, S3B). In wildtype and *miR-34a^−/−^; miR-449^−/−^* DKO choroid plexus explants, basal Shh signaling exhibits a steady decline from E14.5 to P14, yet a robust response to exogenous Shh only occurs in E14.5 and P0, but not in P14 (Sup Fig. S3A, S3B). This developmental decline in Shh responsiveness, together with a developmental decrease in Shh expression (S3D, S3E), results in attenuated Shh signaling that is important to relieve the inhibition of Aqp1 and Atp1a2 expression and promote CSF production during this critical stage for brain ventricle expansion. In *miR-34/449* TKO choroid plexus explants, however, increased cilia number and elongated axonemes drive aberrantly elevated basal Shh signaling level from E14.5 to P14 (Sup Fig. S3C). Interestingly, *miR-34/449* TKO choroid plexus explants fail to respond to exogenous Shh treatment at all developmental stages, although they are capable of responding to Shh when pretreated with Smo inhibitor (Fig. 3D). These findings suggest that *miR-34/449* deficiency markedly enhances the ciliary responsiveness to Shh signaling, such that explant tissue derived basal Shh signaling is sufficient to elicit a maximal Shh response.

To examine the effect of *miR-34/449* on CPEC Shh signaling *in vivo,* we compared *Gli1* expression and ciliary Smo translocation in wildtype, *miR-34a^−/−^; miR-449^−/−^* DKO, and *miR-34/449* TKO choroid plexus tissues from E14.5 to P14 (Fig. 3E-3F). WT and *miR-34/449* DKO choroid plexus tissue exhibit a strong *Gli1* expression at E14.5, which progressively declines by P14 (Fig. 3E). This developmental decline likely reflects the combined effect of decreasing Shh expression and attenuated ciliary responsiveness to Shh signaling (Fig. 3C, Sup Fig.S3D, S3E). Although *miR-34/449* TKO choroid plexus also exhibits a developmental decline of Shh signaling, it exhibits an elevated *Gli1* expression and increased ciliary Smo translocation across all developmental stages examined (Fig. 3E, 3F, Sup Fig. S3A-S3C). Hence, the increased ciliary number and elongated cilia caused by *miR-34/449* deficiency likely enhance their responsiveness to tissue-derived Shh signaling, giving rise to reduced water channel and ion transporter expression and decreased CSF production (Fig. 1F, 1G, 1H).

We then tested whether the aberrant Shh signaling in *miR-34/449* TKO choroid plexus contributes to the widespread reduction in water channel and ion transport expression, particularly Aqp1, the key water channel for CSF production (6). *miR-34/449* TKO choroid plexus explants exhibited reduced apical Aqp1 expression (Fig. 3G), consistent with the decreased Aqp1 expression observed *in vivo* (Fig. 1G, 1H). Importantly, Smo inhibition with KAAD-cyclopamine restored apical Aqp1 expression in *miR-34/449* TKO explants to levels comparable to those of wildtype and *miR-34a^−/−^; miR-449^−/−^* DKO explants (Fig. 3G), suggesting that aberrant Shh signaling underlies the dysregulated Aqp1 expression. These results indicate that *miR-34/449* deficiency in cilia aberrantly elevates Shh signaling, leading to persistent repression of Aqp1 in *miR-34/449* TKO CPECs. Hence, impaired ciliogenesis in *miR-34/449* TKO choroid plexus disrupts the developmental dynamics of Shh signaling required for the rapid CSF production and ventricular expansion during embryonic and postnatal development.

### *miR-34/449* downregulates *Gmnc* to regulate basal body amplification and assembly

*miR-34/449* deficiency produces pleiotropic phenotypes in a variety of multiciliated cell types, including CPECs and airway MCCs. Yet the fundamental defects across all cell types are the excessive basal body amplification and aberrant basal body docking. In CPECs, such defects increase cilia number, disrupt developmentally regulated ciliary shortening, elevate and prolong Sonic hedgehog (Shh) signaling, and consequently, impair cerebrospinal fluid (CSF) production.

To identify the molecular mechanisms underlying the role of *miR-34/449* miRNAs in choroid plexus ciliogenesis, we compared the transcriptomes of *miR-34a*^⁻/⁻^; *miR-449*^⁻/⁻^ DKO and *miR-34/449* TKO choroid plexus at postnatal day 0 (P0) and P14 using RNA-seq (Fig. S4A). The KEGG pathway analysis identifies a significant enrichment of cilia genes among the genes elevated in *miR-34/449* TKO choroid plexus (Fig. 4A, Fig. S4A, Table S1). Real time PCR analyses confirmed a significant upregulation of several multiciliogenesis genes in *miR-34/449* TKO choroid plexus, including transcriptional regulators for multiciliogenesis (*Gmnc*, *Mcidas*, *FoxJ1*) and basal body assembly genes (*Stil*, *Plk4*, *Cep152*) (Fig. 4B, Sup Fig. S4A).These findings are consistent with the increased basal body number and elongated ciliary axonemes observed in *miR-34/449* TKO choroid plexus, because ectopic expression of Gmnc, Mcidas, and FoxJ1 promotes basal body amplification, increases cilia number, and elongates ciliary axonemes (49–51), whereas overexpression of Stil, Plk4, and Cep152 promotes centriole assembly and duplication (52, 53).

**Figure 4.**
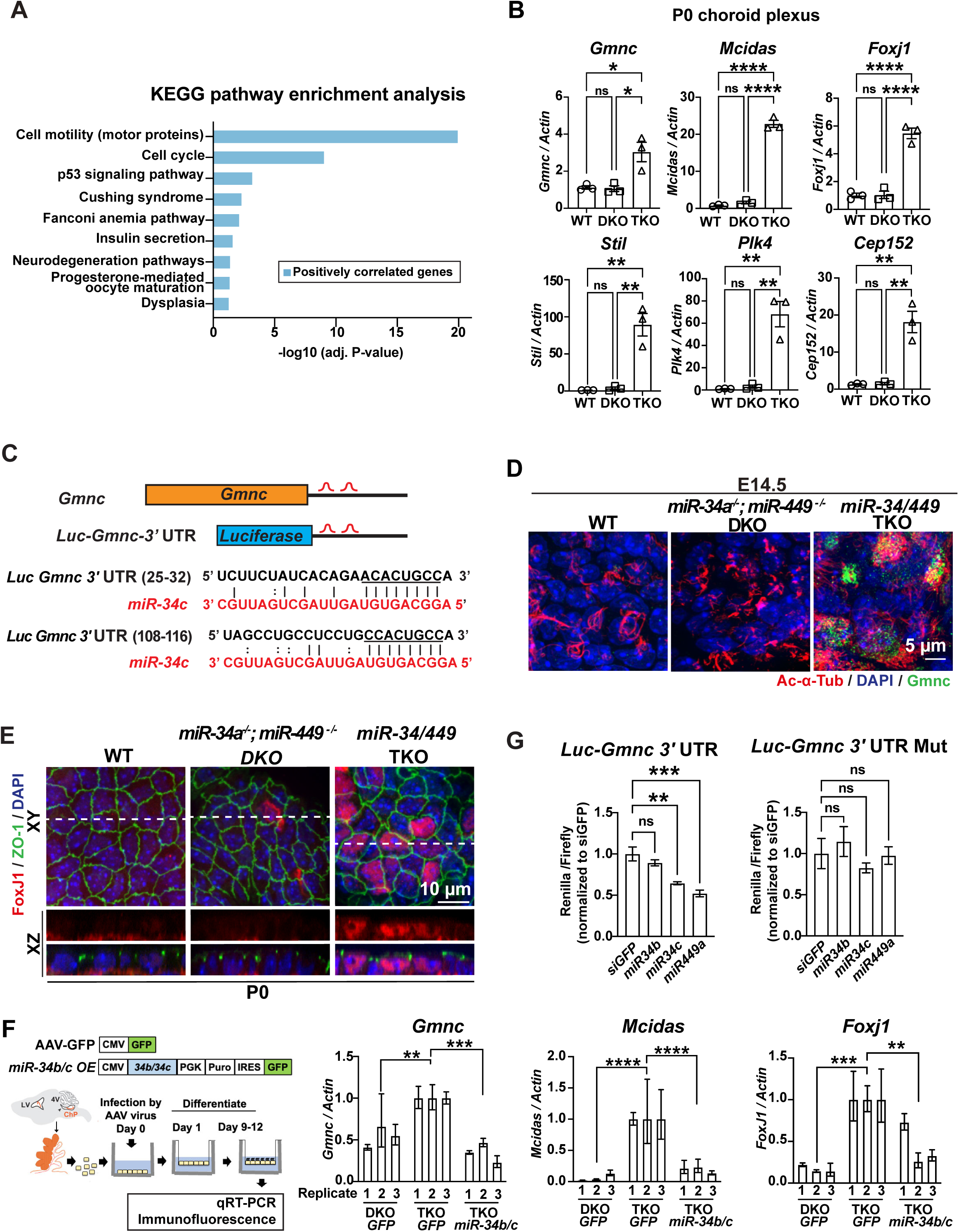
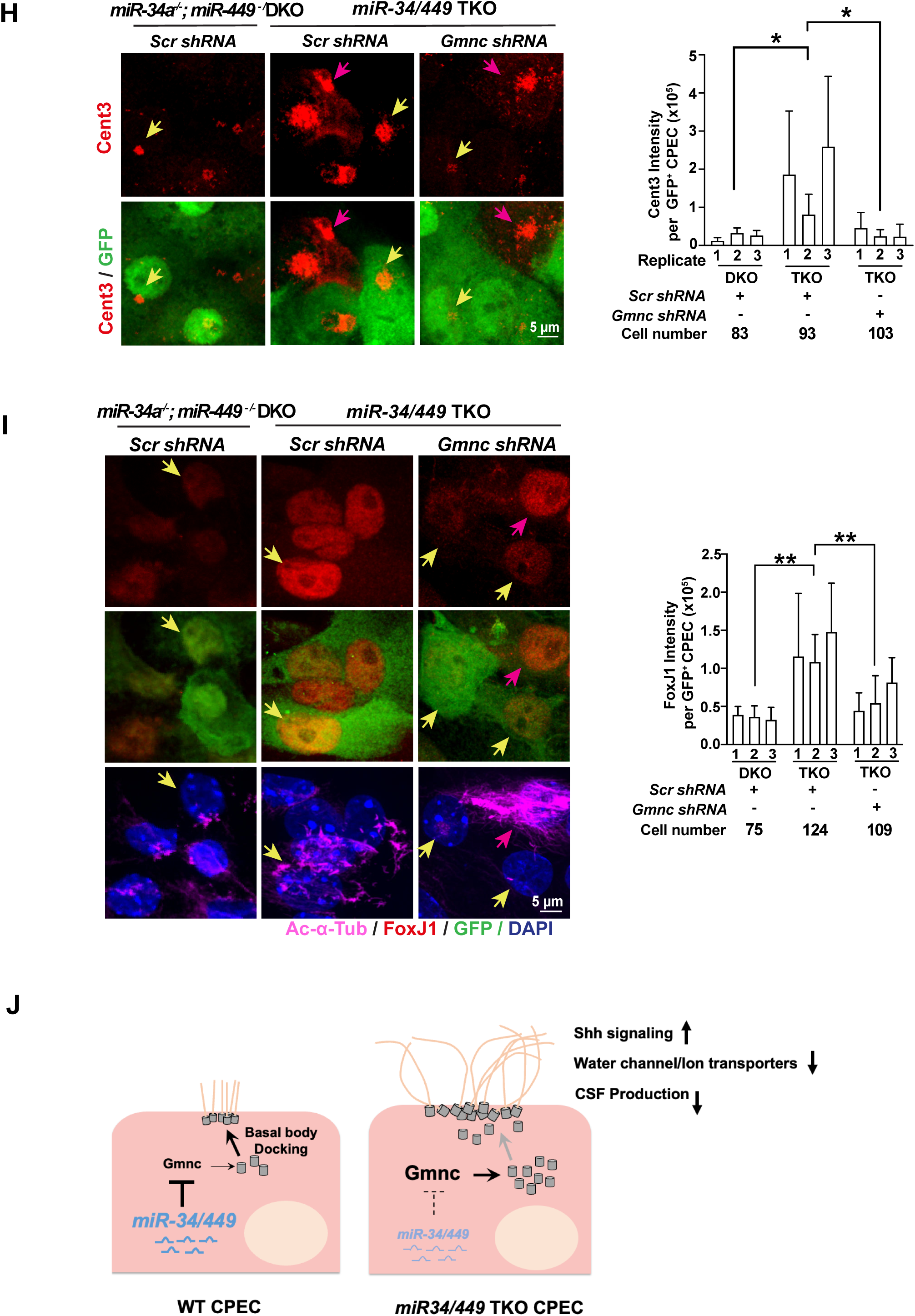
*miR-34/449 miRNAs* target *Gmnc* to regulate basal body assembly and ciliogenesis in CPECs. **A.** RNA-seq analyses were performed to compare the transcriptomes of *miR-34a^−/−^; miR-449^−/−^* DKO and *miR-34/449* TKO choroid plexus at P0 and P14. Differentially upregulated genes (n = 2508) in *miR-34/449* TKO choroid plexus were subjected to KEGG pathway enrichment analyses, revealing significant enrichment of cell motility genes. **B.** Real time PCR analyses validated the upregulation of multiple ciliogenesis genes in P0 TKO choroid plexus, including motile ciliogenesis (*Gmnc, Mcidas, FoxJ1*) and basal body assembly (*Stil, Plk4, Cep152*) genes. n=3 for each genotype group; nested one-way ANOVA and Sidak multiple comparisons tests. Error bars, sem. **C.** The schematic image shows two Targetscan predicted binding sites of *miR-34/449 miRNAs* in the mouse *Gmnc 3’UTR.* **D.** Immunostaining characterized the expression of Gmnc protein in WT, *miR-34a^−/−^; miR-449^−/−^* DKO and *miR-34/449* TKO choroid plexus, revealing an increase of Gmnc nuclear expression in *miR-34/449* TKO CPECs. Quantitation of Gmnc expression is in Sup Fig S4B. **E.** *miR-34/449* deficiency increases the nuclear expression of FoxJ1 in P0 CPECs explant as shown by immunostaining. Representative immunostaining images of FoxJ1 were shown (n=3). Quantitation of FoxJ1 expression is in Sup Fig S4C. **F.** *miR-34/449* overexpression in *miR-34/449* TKO choroid plexus represses the Gmnc/FoxJ1/Multicilin pathway. Left, P0 *miR-34a^−/−^; miR-449^−/−^* DKO and *miR-34/449* TKO monolayer CPEC cultures were infected with AAV2 vector that overexpresses GFP only or overexpresses *mir-34b/34c* and a GFP reporter. The expression of *Gmnc, Mcidas* and *FoxJ1* were assays after 9 days of culture using real time PCR. *Gmnc*: ** *P* = 5.651,t = 5.143, df = 6; *miR-34/449* TKO GFP vs. *miR-34/449* TKO *mir-34b/c*, *** *P* = 0.0004, t = 8.001, df = 6; *Mcidas*: D*miR-449^−/−^* DKO GFP vs. *miR-34/449* TKO GFP, **** *P* < 0.0001, t = 25.51, df = 6; *miR-34/449* TKO GFP vs. *miR-34/449* TKO *mir-34b/c*, **** *P* < 0.0001, t = 21.99, df = 6; *FoxJ1*: *miR-34a^−/−^; miR-449^−/−^* DKO GFP vs. *miR-34/449* TKO GFP, *** *P* = 0.0009, t = 6.854, df = 6; *miR-34/449* TKO GFP vs. *miR-34/449* TKO *mir-34b/c*, ** *P* = 0.0071, t = 34.636, df = 6; n=3; Error bars, sd; one-way ANOVA and Sidak multiple comparisons tests. **G.** Luciferase reporter analyses revealed a *miR-34/449*-dependent repression of *Gmnc.* A luciferase reporter containing a wild-type Gmnc 3’UTR with two *miR-34/449* binding sites can be downregulated by overexpression of *miR-34c* or *miR-449a* mimics in HEK cells. Mutation of the two *miR-34/449* binding sites in this reporter abolished *miR-34/449-*dependent repression. For *Luc*-*Gmnc*-3’UTR, *siGFP* vs. *miR-34b*, n.s., *P* = 0.4358, t = 1.43, df = 8; *siGFP* vs. *miR-34c*, ** *P* = 0.0034, t = 4.929, df = 8; *siGFP* vs. *miR-449a*, *** *P* = 0.0005, t = 6.649, df = 8; For Luc-G*mnc*-3’UTR-Mut, *siGFP* vs. *miR-34b*, n.s., *P* = 0.8671, t = 0.7241, df = 8; *siGFP* vs. *miR-34c*, n.s., *P* = 0.797, t = 0.8649, df = 8; *siGFP* vs. *miR-449a*, n.s., *P* = 0.9992, t = 0.1194, df = 8; n=3, one-way ANOVA and Sidak multiple comparisons tests, error bars, sem. **H, I.** *Gmnc* knockdown in *miR-34/449* TKO CPECs partially rescues basal body assembly defects and restores the appropriate FoxJ1 expression level. P0 *miR-34a^−/−^; miR-449^−/−^* DKO and *miR-34/449* TKO CPEC monolayer cultures were infected by lentiviral vectors expressing either a control shRNA or a *Gmnc* shRNA together with a GFP marker, followed by immunostaining of Centrin-3 (Cent3, basal body) (**H**) or FoxJ1 (**I**), Representative images and the quantification of Cent3 (**H**) or FoxJ1 (**I**) immunostaining intensity were shown in infected GFP^+^ CPECs (yellow arrows) and uninfected GFP^−^ CPEPs (magenta arrows). **H.** *miR-34a^−/−^; miR-449^−/−^* DKO *Scr shRN*A vs. *miR-34/449* TKO *Scr shRN*A, * *P* = 0.0335, t = 3.605, df = 6; *miR-34/449* TKO scramble *shRN*A vs. *miR-34/449* TKO *Gmnc shRN*A, \**P* = 0.0403, t = 3.452, df = 6; nested one-way ANOVA and Sidak multiple comparisons tests. Error bars, sd. **I.** *miR-34a^−/−^; miR-449^−/−^* DKO *Scr shRN*A vs. *miR-34/449* TKO *Scr shRN*A, \*\**P* = 0.0017, t = 6.644, df = 6; *miR-34/449* TKO *Scr shRN*A vs. *miR-34/449* TKO *Gmnc shRN*A, \*\**P* = 0.0087, t = 4.826, df = 6; n=3; error bars, s.d; nested one-way ANOVA and Sidak multiple comparisons tests. **J**. A schematic model illustrating the proposed mechanism by which *miR-34/449* deficiency disrupts choroid plexus multiciliogenesis and signaling. Loss of *miR-34/449* miRNAs causes aberrant basal body amplification, impairs basal body docking, disrupts developmentally regulated ciliary shortening, and results in increased basal body number, cilia number, and axoneme length in CPECs. These ciliary abnormalities in *miR-34/449* deficient CPECs lead to aberrant activation of Sonic Hedgehog (Shh) signaling, reduced expression of water channels and ion transporters required for cerebrospinal fluid (CSF) secretion, and ultimately decreased CSF production.

To identify *miR-34/449* targets, we intersected genes upregulated in *miR-34/449* TKO CPECs with predicted *miR-34/449* targets identified by TargetScan software (Version: Targetscan Release 8.0) (29, 54), and identified 197 candidate targets (Table S2). Among these, *Gmnc* emerges as a strong candidate, as it shows a ∼3 fold upregulation in *miR-34/449* deficient choroid plexus (Fig. 4B), and its 3′ untranslated region (3’UTR) contains two predicted *miR-34/449* binding sites (Fig. 4C).

Gmnc is a key transcriptional regulator for multiciliogenesis, acting downstream of Notch inhibition and upstream of the FoxJ1/Mcidas axis in the developmental hierarchy that specifies multiciliated cell fate and promotes the multiciliogenesis program (50, 55–57). In E14.5 choroid plexus, one of the earliest molecular phenotypes caused by *miR-34/449* deficiency is the marked elevation of *Gmnc* mRNA and protein levels (Fig. 4B, 4D, Sup Fig. S4B). Consistent with FoxJ1 being the key transcriptional target of Gmnc, we also observed an elevated FoxJ1 mRNA and protein level in *miR-34/449* TKO choroid plexus (Fig. 4B, 4E, Sup Fig. S4C). In parallel, we established a CPEC monolayer culture system that enabled *miR-34/449* overexpression using an associated-adenovirus (AAV2) vector (58–60). Single cell suspensions from *miR-34/449* TKO choroid plexus tissue were cultured as monolayers and infected by an AAV vector expressing *miR-34b/c*. *miR-34b/c* overexpression leads to a significant downregulation of *Gmnc*, as well as its downstream targets *FoxJ1* and *Mcidas* (Fig. 4F). These data suggest that Gmnc is subjected to a *miR-34/449-*dependent regulation in CPECs.

Consistent with these findings, a luciferase reporter fused to the *Gmnc* 3′UTR exhibited *miR-34/449*-dependent repression, and this regulation was abolished upon mutation of the two predicted *miR-34/449* binding sites within the 3′UTR (Fig. 4G). Together, these findings suggest that *Gmnc* is likely a direct target of *miR-34/449* miRNAs, subjecting to a *miR-34/449*-dependent regulation.

The elevation of the *Gmnc-FoxJ1/Mcidas* axis in *miR-34/449* TKO choroid plexus could orchestrate a transcriptional program that promotes excessive basal body amplification, as well as aberrant ciliary axoneme outgrowth and assembly (49, 51, 61). To functionally validate Gmnc as a *miR-34/449* target, we cultured *miR-34/449* TKO CPECs as adherent monolayers, and examined the effect of *Gmnc* knockdown on choroid plexus ciliogenesis. Using a lentiviral vector expressing *Gmnc* shRNA and GFP, we achieved efficient Gmnc knockdown as confirmed by real time PCR analyses (Sup Fig. S4D). In this experimental system, we achieved 60-70% lentiviral infection efficiency, and identified *miR-34/449* TKO CPECs with effective *Gmnc* knockdown by GFP expression (Fig. 4H, 4I). When comparing *Gmnc* knockdown *miR-34/449* TKO CPECs with control infected or non-infected *miR-34/449* TKO cells, we observed a marked reduction in basal body numbers (as shown by Centrin 3 immunostaining), ciliary length (as shown by Ac-α-Tub immunostaining) and FoxJ1 expression to levels comparable to *mir-34a^−/^-; mir-449^−/−^* DKO controls (Fig.4H, 4I), Hence, *Gmnc* knockdown rescued the aberrant FoxJ1 elevation, excessive basal body duplication, and elongated ciliary axonemes in *miR-34/449* TKO CPECs, suggesting that *miR-34/449* miRNAs restrain cilia number and ciliary length, at least in part, by modulating Gmnc levels and finetuning the FoxJ1- and Mcidaas-dependent transcriptional program during choroid plexus multiciliogenesis.

## Discussion

Choroid plexus multicilia undergo a unique, yet poorly understood, ciliogenesis program, characterized by a modest cilia number and a developmental decrease in ciliary length. Choroid plexus sensory multicilia and airway motile multicilia share key ciliogenesis programs, as typical mouse ciliogenesis mutants reduce cilia number and length in both cell types (18, 22, 40, 62–64). However, choroid plexus and airway epithelia also exhibit cell type specific cilia numbers and ciliary outgrowth dynamics, suggesting that cell type specific programs also govern aspects of multi-ciliogenesis (18, 25, 65). *miR-34/449* deficient mice are the first model that shows distinct ciliary defects in sensory and motile multicilia: they exhibit fewer and shorter cilia in airway MCCs (33), but more and longer cilia in CPECs (Fig. 1D). Yet both *miR-34/449* deficient cell types share the same underlying cellular defects, excessive basal body amplification and defective basal body docking. Our study provides molecular and functional evidence that *miR-34/449* miRNAs restrict both basal body amplification and ciliary length by suppressing Gmnc, an upstream activator of the Mcidas/FoxJ1 transcriptional cascade that governs ciliogenesis. A modest upregulation of Gmnc in choroid plexus, such as that induced by *miR-34/449* deficiency, triggers a considerable elevation of its downstream targets that collectively cause excessive basal body amplification.

Why does *miR-34/449* deficiency cause seemingly contrasting ciliary phenotypes in airway MCCs and CPECs? In airway MCCs, basal bodies normally dock uniformly across the entire apical surface, leaving no apical surface available for additional basal body docking.

Consequently, decreased docking efficiency caused by *miR-34/449* deficiency reduces cilia number and ciliary length despite the increased basal body amplification. In contrast, basal body docking in CPECs is normally confined to a central apical region. Excessive basal body amplification in *miR-34/449* deficient CPECs promotes ectopic docking across a broader apical region, ultimately increasing cilia number despite impaired docking efficiency. Interestingly, *miR-34/449* deficiency does not impair choroid plexus ciliary length initially, but the resulting docking defects likely disrupt the developmental decrease of ciliary length, thereby resulting in longer and dispersed cilia in mature choroid plexus epithelia, which failed to undergo ciliary shortening. Together, these findings position m*iR-34/449* miRNAs as context-dependent regulators that tune ciliogenic programs to tissue-specific developmental trajectories.

In mouse embryos, choroid plexus cilia transduce a non-canonical Sonic hedgehog (Shh) signaling pathway that downregulates the expression of ion channels and water transporters, thereby reducing cerebrospinal fluid (CSF) production to maintain CSF homeostasis (Mao et al., 2025). During postnatal development, choroid plexus cilia undergo a progressive decrease in ciliary length and exhibit an attenuated response to Shh signaling. Reduced ciliary sensing capacity, together with decreased levels of Shh ligand, contributes to attenuation of Shh signaling, derepression of water channel and ion transporter expression, and a developmental increase in CSF production that supports rapid expansion of the brain ventricles. The loss of *miR-34/449* miRNAs disrupts these developmental dynamics of Shh signaling, leading to sustained elevation of Shh signaling throughout development. Consequently, persistent repression of water channel and ion transporter expression reduces CSF production and ultimately promotes microcephaly.

Taken together, *miR-34/449* miRNAs not only promote multiciliogenesis in CPECs, but also enable dynamic modulation of Shh signaling and CSF production in development, through the regulation of ciliary length. These findings align with emerging evidence that cilia turnover, like ciliogenesis, is an actively regulated genetic program rather than a passive process. Emerging evidence suggests that different ciliated cells employ distinct mechanisms to remodel ciliary length and ciliary function as a part of physiological functions. Our studies identified *miR-34/449* miRNAs as important regulators for basal body dynamics, and the proper basal body docking is likely a prerequisite for continuous, developmentally cilia shortening in choroid plexus. A recent genome-wide CRISPR activation screen, however, identified a signaling cascade actively driving primary cilia disassembly (66). The authors found that SARM1 acts downstream of F2R G protein-coupled receptor acts and engages ryanodine receptor-mediated peri-centrosomal calcium signaling to activate RhoA, together forming a pathway necessary and sufficient for cilia disassembly in focal cortical dysplasia, a neurodevelopmental disorder marked by intractable epilepsy (*Elliott et al*., 2025). Altogether, our findings highlight dynamic, rather than static, ciliary functions as important regulators of physiology across diverse ciliated cell types.

## Author Contributions

S.M., R.S. G.U. and L.H. are the main contributors to this study. R.S., S.M. and L.H. conceptualized the study, formulated the hypothesis and designed the key experiments. R.S initiated the project and identified aberrant choroid plexus ciliogenesis and microcephaly in *miR-34/449* TKO mice, and characterized this phenotype using immunostaining, SEM, TEM (in collaboration with D.J), and FIB-SEM (in collaboration with S.P., C.S.X., S.J., A.Z. H.H. and S.U.). S.J. and Z.X. performed RNA-seq experiments to identify *miR-34/449* targets. S.M., with the help of A.J. and D.L., made substantial contributions to the understanding of aberrant Shh signaling caused by cilia defects in *miR-34/449* deficient choroid plexus. S.M., R.S. S.J. developed the hypothesis of Gmnc being the key *miR-34/449* targets, and S.M. performed most experiments for target validation. M.F.W. and S.M. performed MRI on CSF measurement. S.M., R.S and S.J developed the CPEC monolayer culture system, and S.M. employed this experimental system to functionally validate Gmnc as a key *miR-34/449* target. S.M., and L.H. are the key contributors to the writing of the manuscript.

## Acknowledgments

We thank W Chen, J Nealon, P Lu, M. Kinisu, T Machen, P Lishko, D Bautista, and C Ott, E Monuki, E Vladar and S Liddelow for technical input and stimulating discussion, K McDonald and P Kysar for SEM and TEM analyses, S Brody for *Foxj1^−/−^* mice, the UC-Berkeley Electron Microscope Laboratory for assistance in electron microscopy sample preparation and data collection, and M Lehtinen for advice and reagents for the choroid plexus and CSF-related experiments. L.H. is a Thomas and Stacey Siebel Distinguished Chair Professor, supported by a Howard Hughes Medical Institute (HHMI) Faculty Scholar award, a Bakar Fellow award at UC Berkeley, and grants from the National Institutes of Health (1R01GM114414, R01NS120287) and from Nan Fung life sciences. R.S. was supported by a K99 Pathway to independent award by NIH K99HL128912. S.M. was supported by the Postdoctoral Fellowship Award from TRDRP (T30FT1019). S.P. C.S.X. and H.F.H. are supported by the Howard Hughes Medical Institute. S.U. was partially funded by the Philomathia Foundation. S.U. is supported by the Sloan Foundation, and the Chan Zuckerberg Initiative Imaging Scientist program. Z. X. was supported by NIH (1R01NS137584). L.H. and S.U. are supported by the Chan Zuckerberg Biohub – San Francisco Investigator program

## Supplementary table legend

**Table S1.**
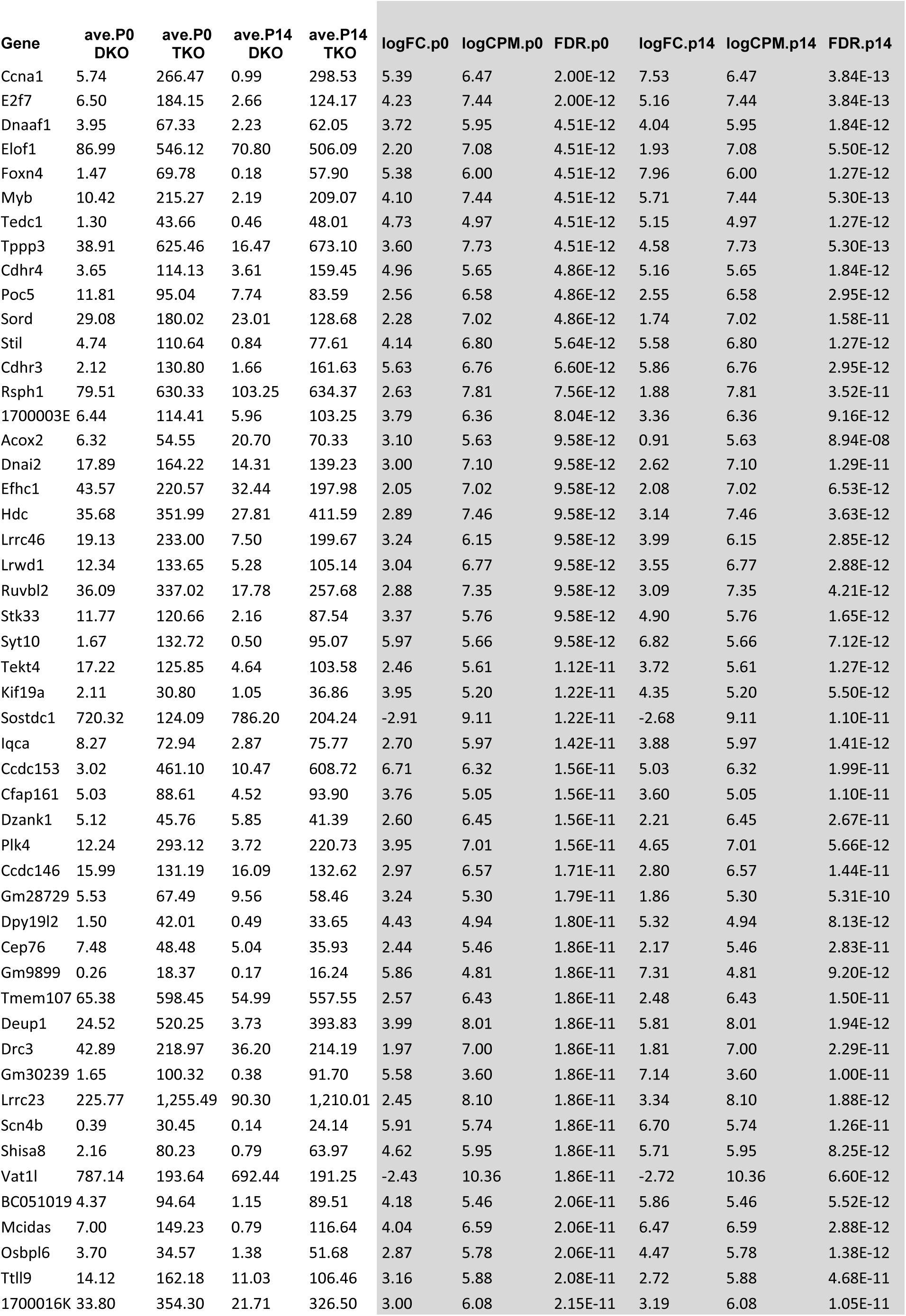
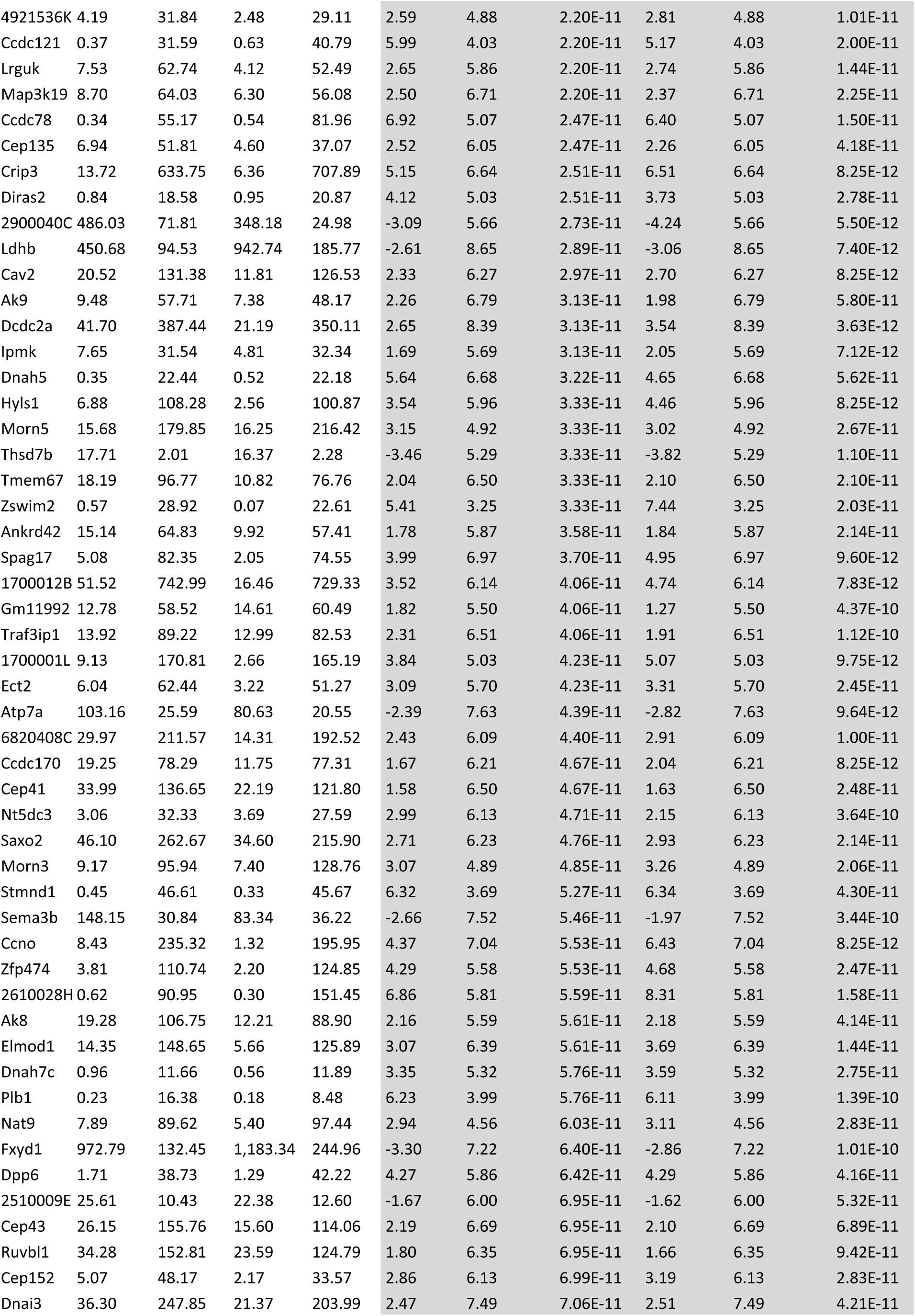
Top 100 dynamically expressed genes in P0 and p14 DKO and TKO choroid plexus, related to Figure 4.

**Table S2.**
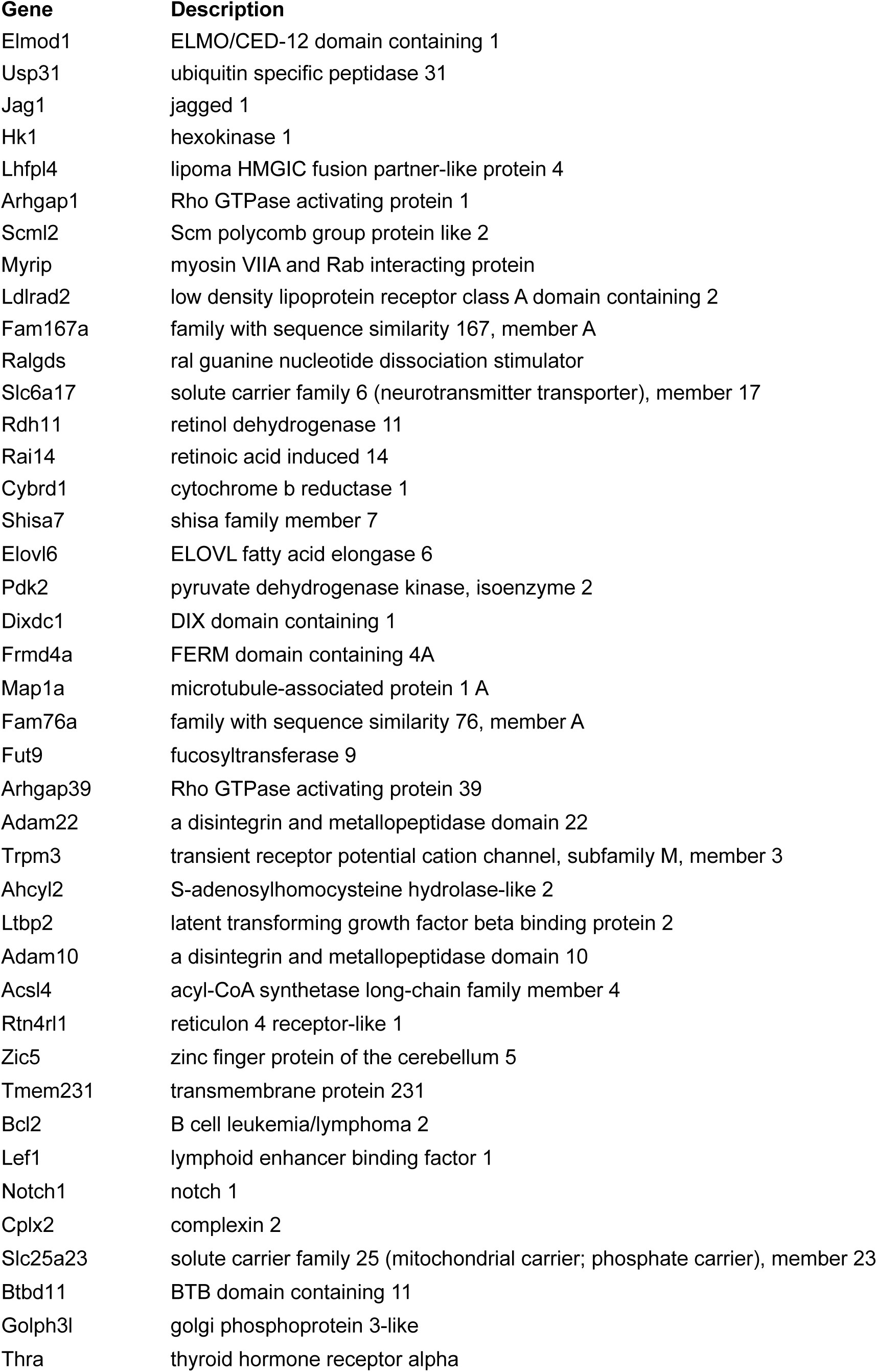
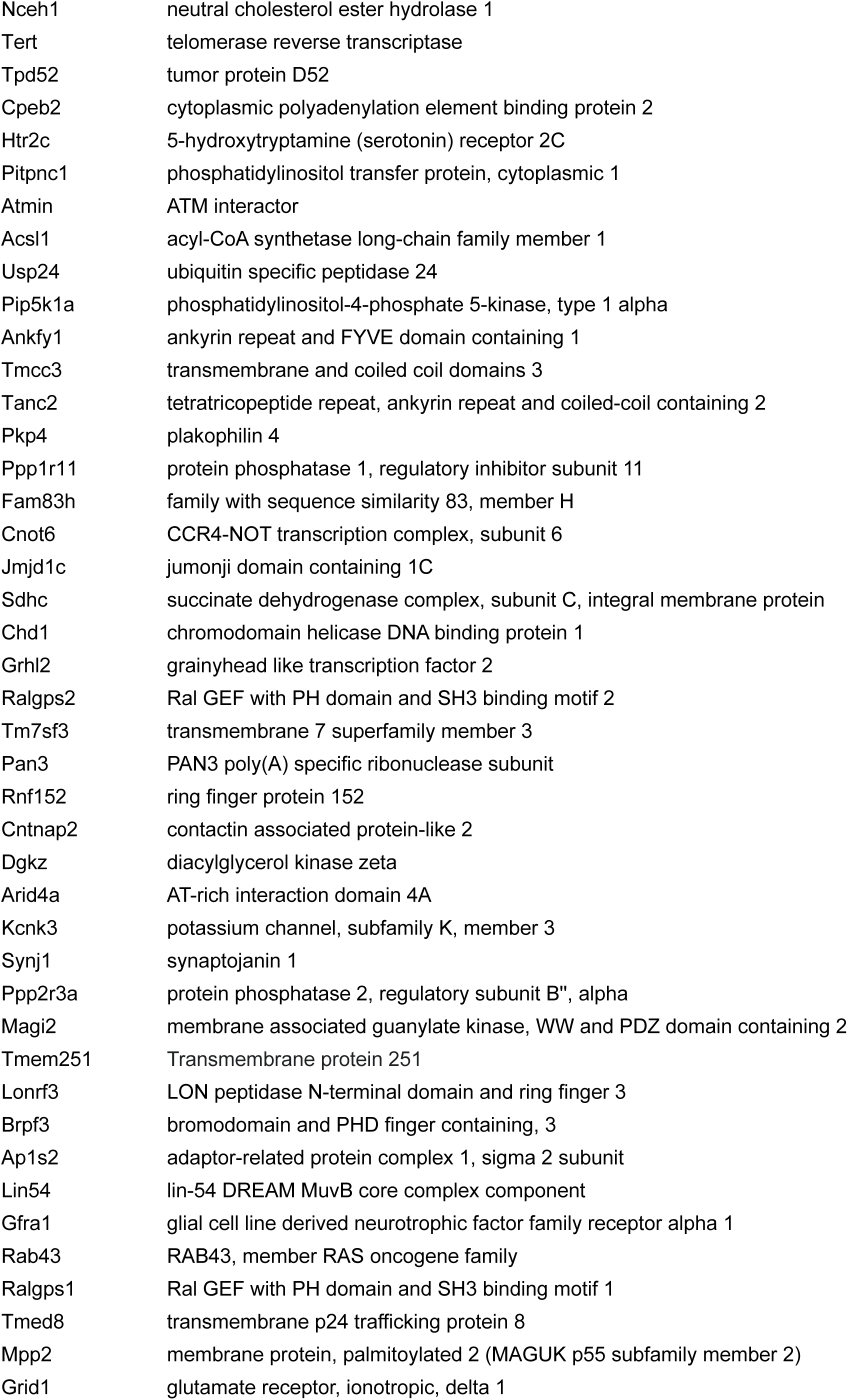
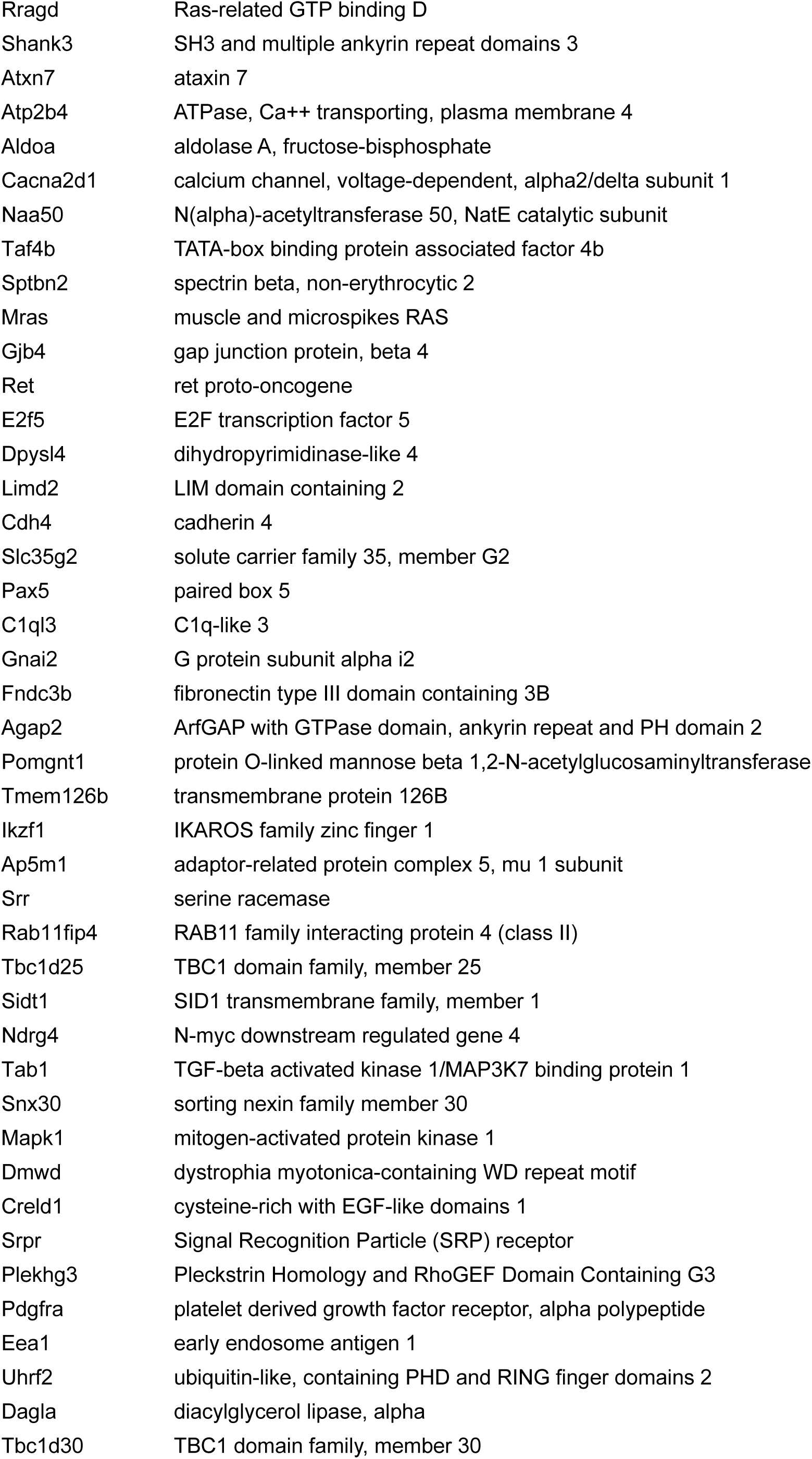
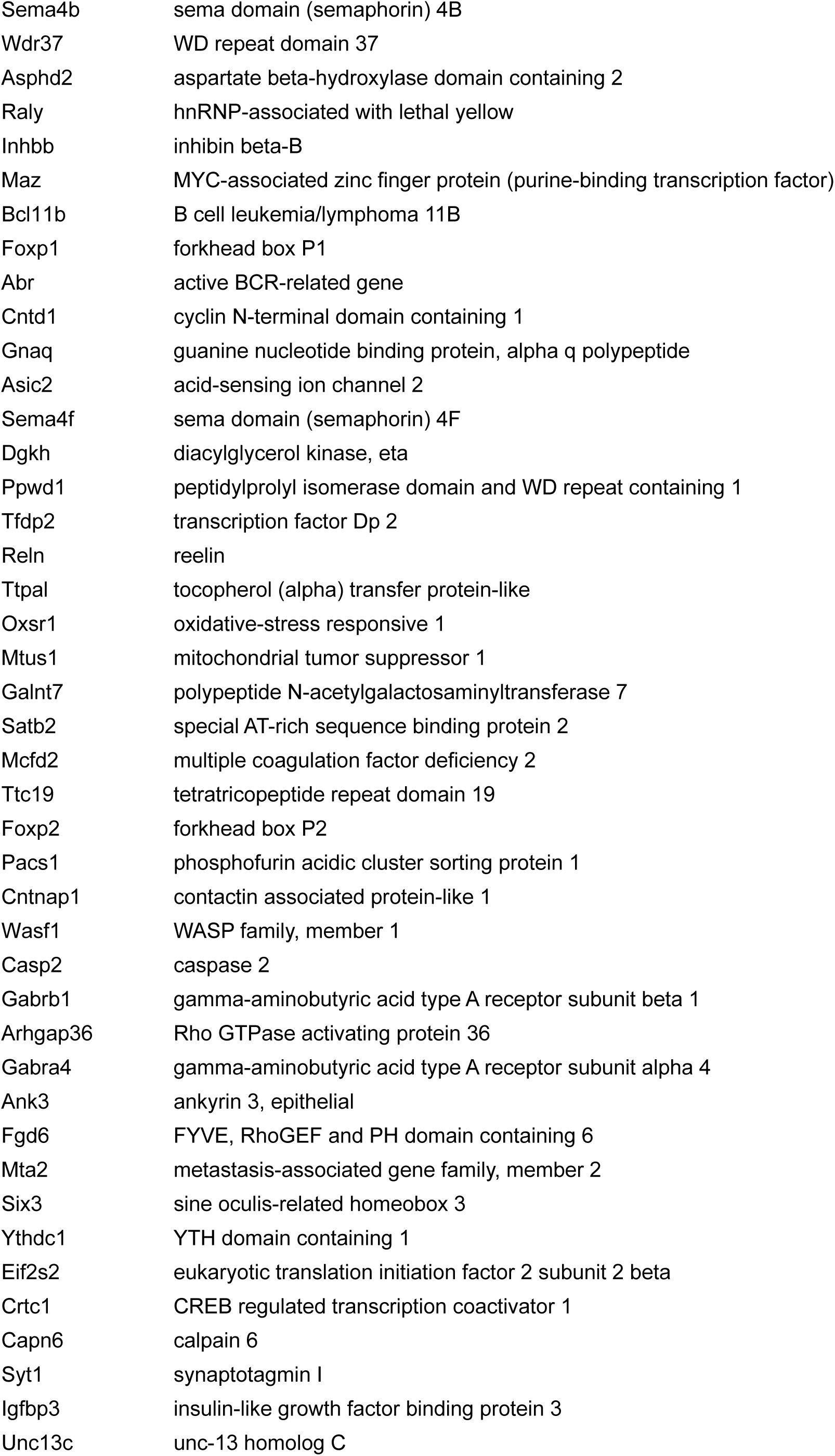
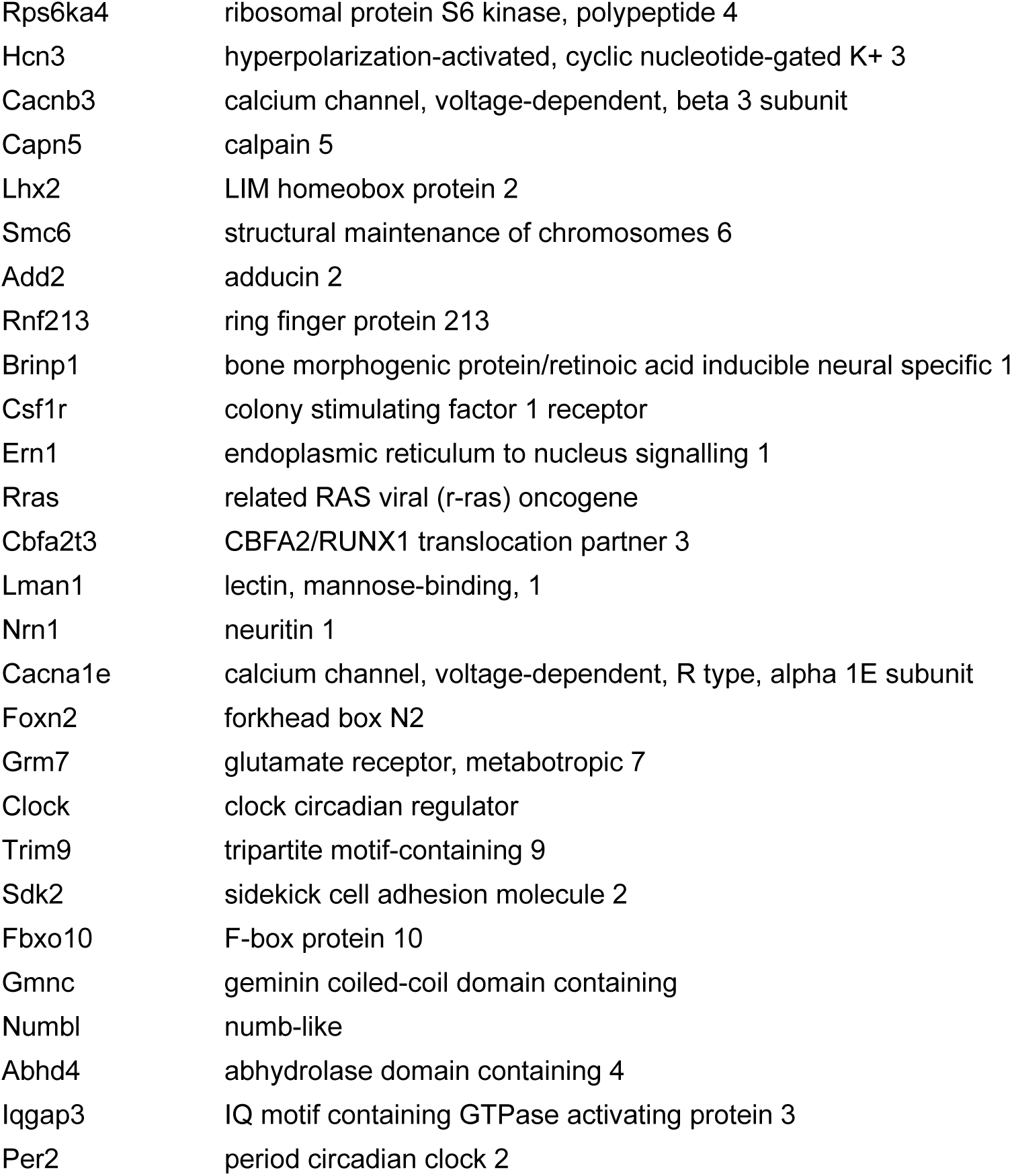
Upregulated genes that are predicted miR-34/449-targets.

**Table S3.** Description for quantitation and statistical tests analysis.

| Figure | Statistics method | Error bar |
| --- | --- | --- |
| **Fig. 1F** | P7 DKO vs. P7 TKO, $n=3$, ***$P = 0.0005$, $t = 10.57$, $df = 4$; unpaired two-tailed Student's t-test | s.e.m. |
| **Fig. 1H** | **Aqp1**, P14 WT vs. P14 DKO, ns, $P = 0.3086$, $t = 1.838$, $df = 6$; P14 WT vs. P14 TKO, *$P = 0.0214$, $t = 3.991$, $df = 6$; P14 DKO vs. P14 TKO, **$P = 0.0032$, $t = 5.884$, $df = 6$.<br>**Atp1a2**, P14 WT vs. P14 DKO, ns, $P = 0.8217$, $t = 0.8322$, $df = 6$; P14 WT vs. P14 TKO, *$P = 0.0472$, $t = 3.321$, $df = 6$; P14 DKO vs. P14 TKO, *$P = 0.0172$, $t = 4.186$, $df = 6$.<br>**Nkcc1**, P14 WT vs. P14 DKO, ns, $P = 0.1004$, $t = 2.72$, $df = 6$; P14 WT vs. P14 TKO, *$P = 0.0366$, $t = 3.53$, $df = 6$; P14 DKO vs. P14 TKO, **$P = 0.0025$, $t = 6.172$, $df = 6$; $n=3$, Nested one-way ANOVA and Sidak multiple comparisons tests.<br>**Slc4a10**, P14 WT vs. P14 DKO, ns, $P = 0.863$, $t = 0.7448$, $df = 6$; P14 WT vs. P14 TKO, *$P = 0.0187$, $t = 4.112$, $df = 6$; P14 DKO vs. P14 TKO, *$P = 0.0483$, $t = 3.302$, $df = 6$.<br>**Ae2**, P14 WT vs. P14 DKO, ns, $P = 0.412$, $t = 1.593$, $df = 6$; P14 WT vs. P14 TKO, **$P = 0.0014$, $t = 6.883$, $df = 6$; P14 DKO vs. P14 TKO, ***$P = 0.0006$, $t = 8.126$, $df = 6$. | s.e.m. |
| **Fig. 2G** | Basal body number, WT vs. DKO, ns, $P = 0.9958$, $t = 0.2042$, $df = 48$; WT vs. TKO, ****$P < 0.0001$, $t = 21.47$, $df = 48$; DKO vs. TKO, ****$P < 0.0001$, $t = 19.44$, $df = 48$; one-way ANOVA and Sidak multiple comparisons tests. | s.d. |
| **Fig. 3A** | **For WT**, DMSO vs. Shh, ****$P < 0.0001$, $t = 10.52$, $df = 8$; WT DMSO vs. KAAD, ns, $P = 0.0806$, $t = 2.687$, $df = 8$; WT DMSO vs. Shh + KAAD, ns, $P = 0.9981$, $t = 0.1618$, $df = 8$.<br>**For DKO**, DMSO vs. Shh, *$P = 0.0182$, $t = 3.693$, $df = 8$; DKO DMSO vs. KAAD, ns, $P = 0.2766$, $t = 1.845$, $df = 8$; DKO DMSO vs. Shh + KAAD, ns, $P = 0.8200$, $t = 0.8211$, $df = 8$.<br>**For TKO**, DMSO vs. Shh, ns, $P = 0.1568$, $t = 2.242$, $df = 8$; TKO DMSO vs. KAAD, **$P = 0.0013$, $t = 5.744$, $df = 8$; TKO DMSO vs. Shh + KAAD, ***$P = 0.0004$, $t = 6.727$, $df = 8$; one-way ANOVA and Sidak multiple comparisons tests. | s.d. |
| **Fig. 3C** | P14 WT PBS vs. WT + Shh, ns, $P = 0.11$, $t = 2.047$, $df = 4$. P14 DKO PBS vs. DKO + Shh, ns, $P = 0.302$, $t = 1.184$, $df = 4$. P14 | s.e.m. |
| | TKO PBS vs. TKO + Shh, ns, $P = 0.1066$, $t = 2.075$, $df = 4$; $n = 3$, unpaired two-tailed Student's t-test. | |
| **Fig. 3D** | Ciliary Smo intensity, DKO WT PBS ($n=54$) vs. DKO Shh ($n=61$), ***$P = 0.0001$, $t = 3.961$, $df = 113$; TKO WT PBS ($n=39$) vs. TKO Shh ($n=65$), ****$P < 0.0001$, $t = 5.8$, $df = 102$; unpaired two-tailed Student's t-test. | s.d. |
| **Fig. 3E** | For E14.5 *Gli1*, E14.5 WT vs. E14.5 DKO, ns, $P = 0.9953$, $t = 0.2214$, $df = 6$; E14.5 WT vs. E14.5 TKO, ns, $P = 0.0161$, $t = 4.248$, $df = 6$; E14.5 DKO vs. E14.5 TKO, *, $P = 0.0127$, $t = 4.469$, $df = 6$.<br>For P0 *Gli1*, P0 WT vs. P0 DKO, ns, $P = 0.9905$, $t = 0.2808$, $df = 6$; P0 WT vs. P0 TKO, **$P = 0.0017$, $t = 6.616$, $df = 6$; P0 DKO vs. P0 TKO, ** $P = 0.0022$, $t = 6.335$, $df = 6$.<br>For P14 *Gli1*, P14 WT vs. P14 DKO, ns, $P = 0.9499$, $t = 0.5052$, $df = 6$; P14 WT vs. P14 TKO, *$P = 0.0111$, $t = 4.593$, $df = 6$; P14 DKO vs. P14 TKO, *$P = 0.0192$, $t = 4.088$, $df = 6$; $n=3$, one-way ANOVA and Sidak multiple comparisons tests. | s.e.m. |
| **Fig. 3G** | Quantitation of apical Aqp1 expression in ciliated CPECs, DKO DMSO vs. TKO DMSO, **$P = 0.0095$, $t = 4.154$, $df = 8$; DKO DMSO vs. KAAD, *$P = 0.019$, $t = 4.154$, $df = 8$; TKO DMSO vs. KAAD, ** $P = 0.0099$, $t = 4.648$, $df = 8$; DKO KAAD vs. TKO KAAD, ns, $P = 0.1041$, $t = 2.959$, $df = 8$; nested unpaired One-way ANOVA and Sidak multiple comparison. | s.d. |
| **Fig. 4B** | ***Gmnc***, P0 WT vs. P0 DKO, ns, $P = 0.9986$, $t = 0.148$, $df = 6$; P0 WT vs. P0 TKO, * $P = 0.0165$, $t = 4.224$, $df = 6$; P0 DKO vs. P0 TKO, * $P = 0.0141$, $t = 4.372$, $df = 6$.<br>***Mcidas***, P0 WT vs. P0 DKO, ns, $P = 0.7533$, $t = 0.9627$, $df = 6$; P0 WT vs. P0 TKO, **** $P < 0.0001$, $t = 26.3$, $df = 6$; P0 DKO vs. P0 TKO, **** $P < 0.0001$, $t = 25.33$, $df = 6$.<br>***FoxJ1***, P0 WT vs. P0 DKO, ns, $P = 0.9997$, $t = 0.08799$, $df = 6$; P0 WT vs. P0 TKO, **** $P < 0.0001$, $t = 11.13$, $df = 6$; P0 DKO vs. P0 TKO, **** $P < 0.0001$, $t = 11.04$, $df = 6$; one-way ANOVA and Sidak multiple comparisons tests.<br>***Stil***, P0 WT vs. P0 DKO, ns, $P = 0.9964$, $t = 0.2011$, $df = 6$; P0 WT vs. P0 TKO, ** $P = 0.0012$, $t = 7.078$, $df = 6$; P0 DKO vs. P0 TKO, ** $P = 0.0014$, $t = 6.877$, $df = 6$.<br>***Plk4***, P0 WT vs. P0 DKO, ns, $P = 0.9962$, $t = 0.2063$, $df = 6$; P0 WT vs. P0 TKO, ** $P = 0.0011$, $t = 7.186$, $df = 6$; P0 DKO vs. P0 TKO, ** $P = 0.0013$, $t = 6.979$, $df = 6$. | s.e.m. |
| | **Cep152**, P0 WT vs. P0 DKO, ns, $P = 0.9968$, $t = 0.07331$, $df = 6$; P0 WT vs. P0 TKO, ** $P = 0.0012$, $t = 7.093$, $df = 6$; P0 DKO vs. P0 TKO, ** $P = 0.0013$, $t = 7.019$, $df = 6$. | |
| **Fig. 4F** | **Gmnc**, DKO+mock vs. TKO+mock, ** $P = 5.651$, $t = 5.143$, $df = 6$; TKO+mock vs. TKO+*miR34b/c*, *** $P = 0.0004$, $t = 8.001$, $df = 6$. **Mcidas**, DKO+mock vs. TKO+mock, **** $P < 0.0001$, $t = 25.51$, $df = 6$; TKO+mock vs. TKO+*miR34b/c*, **** $P < 0.0001$, $t = 21.99$, $df = 6$. **FoxJ1**, DKO+mock vs. TKO+mock, *** $P = 0.0009$, $t = 6.854$, $df = 6$; TKO+mock vs. TKO+*miR34b/c*, ** $P = 0.0071$, $t = 34.636$, $df = 6$; one-way ANOVA and Sidak multiple comparisons tests. | s.d. |
| **Fig. 4G** | WT *Gmnc*-3'UTR, *siGFP* vs. *miR34b* mimic, ns, $P = 0.4358$, $t = 1.43$, $df = 8$; *siGFP* vs. *miR34c* mimic, ** $P = 0.0034$, $t = 4.929$, $df = 8$; *siGFP* vs. *miR449a* mimic, *** $P = 0.0005$, $t = 6.649$, $df = 8$. For mutated *Gmnc*-3'UTR, *siGFP* vs. *miR34b* mimic, ns, $P = 0.8671$, $t = 0.7241$, $df = 8$; *siGFP* vs. *miR34c* mimic, ns, $P = 0.797$, $t = 0.8649$, $df = 8$; *siGFP* vs. *miR449a* mimic, ns, $P = 0.9992$, $t = 0.1194$, $df = 8$; one-way ANOVA and Sidak multiple comparisons tests. | s.e.m. |
| **Fig. 4H** | **Cent3 intensity**, DKO+*Scr shRNA* vs. TKO+*Scr shRNA*, **$P = 0.0043$, $t = 5.143$, $df = 6$; TKO+*Scr shRNA* vs. TKO+*Gmnc shRNA*, **$P = 0.003$, $t = 5.505$, $df = 6$; nested one-way ANOVA and Sidak multiple comparisons tests. | s.d |
| **Fig. 4I** | **Quantitation of FoxJ1 intensity.**<br>DKO+*Scr shRNA* vs. TKO+*Scr shRNA*, ** $P = 0.0017$, $t = 6.644$, $df = 6$; TKO+*Scr shRNA* vs. TKO+*Gmnc shRNA*, **$P = 0.0087$, $t = 4.826$, $df = 6$; nested one-way ANOVA and Sidak multiple comparisons tests. | s.d |
| **Sup Fig. S1B** | Brain weight quantification, $n=3$; DKO vs. TKO, *$P = 0.0253$, $t = 3.483$, $df = 4$, unpaired two-tailed Student's t-test. | s.e.m. |
| **Sup Fig. S1D** | **Aqp1**, E14.5 WT vs. E14.5 DKO, ns, $P = 0.1147$, $t = 2.616$, $df = 6$; E14.5 WT vs. E14.5 TKO, ns, $P = 0.9919$, $t = 0.2660$, $df = 6$; E14.5 DKO vs. E14.5 TKO, ns, $P = 0.1532$, $t = 2.392$, $df = 6$.<br>**Atp1a2**, E14.5 WT vs. E14.5 DKO, ns, $P = 0.9919$, $t = 0.2663$, $df = 6$; E14.5 WT vs. E14.5 TKO, ns, $P = 0.124$, $t = 2.556$, $df = 6$; E14.5 DKO vs. E14.5 TKO, ns, $P = 0.1634$, $t = 2.341$, $df = 6$.<br>**Slc4a10**, E14.5 WT vs. E14.5 DKO, ns, $P = 0.9904$, $t = 0.2816$, $df = 6$; E14.5 WT vs. E14.5 TKO, ns, $P = 0.9986$, $t = 0.1452$, $df = 6$; E14.5 DKO vs. E14.5 TKO, ns, $P = 0.9989$, $t = 0.1336$, $df = 6$. | s.e.m. |
|  | **Ae2**, E14.5 WT vs. E14.5 DKO, ns, $P = 0.0382$, $t = 3.494$, $df = 6$; E14.5 WT vs. E14.5 TKO, ns, $P = 0.7899$, $t = 0.8945$, $df = 6$; E14.5 DKO vs. E14.5 TKO, ns, $P = 0.1188$, $t = 2.589$, $df = 6$.<br>**Nkcc1**, E14.5 WT vs. E14.5 DKO, ns, $P = 0.2367$, $t = 2.051$, $df = 6$; E14.5 WT vs. E14.5 TKO, ns, $P = 0.9004$, $t = 0.6556$, $df = 6$; E14.5 DKO vs. E14.5 TKO, ns, $P = 0.5262$, $t = 1.368$, $df = 6$; $n=3$, Nested one-way ANOVA and Sidak multiple comparisons tests. |  |
| Sup Fig. S1E | **Aqp1**, E14.5 WT vs. E14.5 DKO, ns, $P = 0.9962$, $t = 0.2057$, $df = 6$; E14.5 WT vs. E14.5 TKO, $*P = 0.0127$, $t = 4.464$, $df = 6$; E14.5 DKO vs. E14.5 TKO, $*P = 0.0103$, $t = 4.669$, $df = 6$.<br>**Aqp1**, P14 WT vs. P14 DKO, ns, $P = 0.3129$, $t = 1.826$, $df = 6$; P14 WT vs. P14 TKO, $*P = 0.0159$, $t = 4.259$, $df = 6$; P14 DKO vs. P14 TKO, $**P = 0.0027$, $t = 6.086$, $df = 6$.<br>**Atp1a2**, E14.5 WT vs. E14.5 DKO, ns, $P = 0.9898$, $t = 0.2874$, $df = 6$; E14.5 WT vs. E14.5 TKO, $*P = 0.0387$, $t = 3.484$, $df = 6$; E14.5 DKO vs. E14.5 TKO, ns, $P = 0.0550$, $t = 3.196$, $df = 6$.<br>**Atp1a2**, P14 WT vs. P14 DKO, ns, $P = 0.6348$, $t = 1.173$, $df = 6$; P14 WT vs. P14 TKO, $**P = 0.0096$, $t = 4.738$, $df = 6$; P14 DKO vs. P14 TKO, $*P = 0.0352$, $t = 3.565$, $df = 6$.<br>**Nkcc1**, E14.5 WT vs. E14.5 DKO, ns, $P = 0.6187$, $t = 1.201$, $df = 6$; E14.5 WT vs. E14.5 TKO, $*P = 0.0127$, $t = 4.468$, $df = 6$; E14.5 DKO vs. E14.5 TKO, $**P = 0.0039$, $t = 5.67$, $df = 6$;<br>**Nkcc1**, P14 WT vs. P14 DKO, ns, $P = 0.4232$, $t = 1.57$, $df = 6$; P14 WT vs. P14 TKO, $****P < 0.0001$, $t = 12.19$, $df = 6$; P14 DKO vs. P14 TKO, $****P < 0.0001$, $t = 13.76$, $df = 6$;<br>**Ae2**, E14.5 WT vs. E14.5 DKO, ns, $P = 0.9978$, $t = 0.1714$, $df = 6$; E14.5 WT vs. E14.5 TKO, ns, $P = 0.07$, $t = 3.003$, $df = 6$; E14.5 DKO vs. E14.5 TKO, ns, $P = 0.087$, $t = 2.832$, $df = 6$;<br>**Ae2**, P14 WT vs. P14 DKO, ns, $P = 0.0552$, $t = 3.193$, $df = 6$; P14 WT vs. P14 TKO, $*P = 0.0107$, $t = 4.632$, $df = 6$; P14 DKO vs. P14 TKO, $***P = 0.0007$, $t = 7.826$, $df = 6$;<br>**Slc4a10**, E14.5 WT vs. E14.5 DKO, ns, $P = 0.6672$, $t = 1.116$, $df = 6$; E14.5 WT vs. E14.5 TKO, $**P = 0.0029$, $t = 5.995$, $df = 6$; E14.5 DKO vs. E14.5 TKO, $**P = 0.0083$, $t = 4.879$, $df = 6$;<br>**Slc4a10**, P14 WT vs. P14 DKO, ns, $P = 0.9746$, $t = 0.3959$, $df = 6$; P14 WT vs. P14 TKO, $**P = 0.0028$, $t = 46.026$, $df = 6$; P14 DKO vs. P14 TKO, $**P = 0.002$, $t = 6.422$, $df = 6$; $n=3$, Nested one-way ANOVA and Sidak multiple comparisons tests. | s.e.m. |
| **Sup Fig. 4B** | **Gmnc intensity**, E14.5 WT vs E14.5 DKO, ns, $P = 0.294$, $t = 1.877$, $df = 6$; E14.5 WT vs E14.5 TKO, *** $P = 0.0006$, $t = 8.105$, $df = 6$; E14.5 DKO vs E14.5 TKO, *** $P = 0.0002$, $t = 9.983$, $df = 6$; one-way ANOVA and Sidak multiple comparisons tests. one-way ANOVA and Sidak multiple comparisons tests. | s.e.m. |
| **Sup Fig. 4C** | **FoxJ1 intensity**, P0 WT vs. P0 DKO, ns, $P = 0.2298$, $t = 1.738$, $df = 280$; P0 WT vs. P0 TKO, **** $P < 0.0001$, $t = 20.56$, $df = 280$; P0 DKO vs. P0 TKO, **** $P < 0.0001$, $t = 23.08$, $df = 280$; one-way ANOVA and Sidak multiple comparisons tests. | s.d. |
| **Sup Fig. 4D** | **Gmnc**, DKO+Scr shRNA vs. TKO+Scr shRNA, **$P = 0.0043$, $t = 5.143$, $df = 6$; TKO+Scr shRNA vs. TKO+Gmnc shRNA, **$P = 0.003$, $t = 5.505$, $df = 6$.<br>**Mcidas**, DKO+Scr shRNA vs. TKO+Scr shRNA, ****$P < 0.0001$, $t = 12$, $df = 6$; TKO+Scr shRNA vs. TKO+Gmnc shRNA, ***$P = 0.0007$, $t = 7.302$, $df = 6$.<br>**FoxJ1**, DKO+Scr shRNA vs. TKO+Scr shRNA, **$P = 0.0038$, $t = 5.256$, $df = 6$; TKO+Scr shRNA vs. TKO+Gmnc shRNA, *$P = 0.0326$, $t = 3.3$, $df = 6$; one-way ANOVA and Sidak multiple comparisons tests. | s.d. |

**Table S4.**
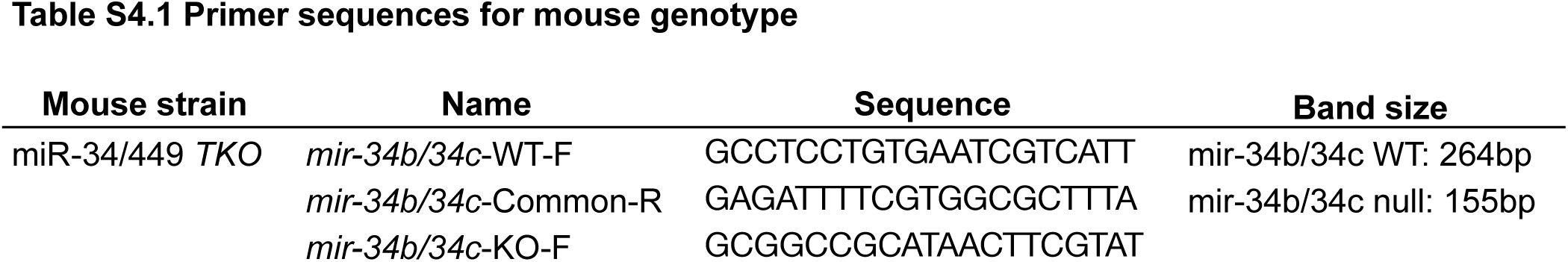
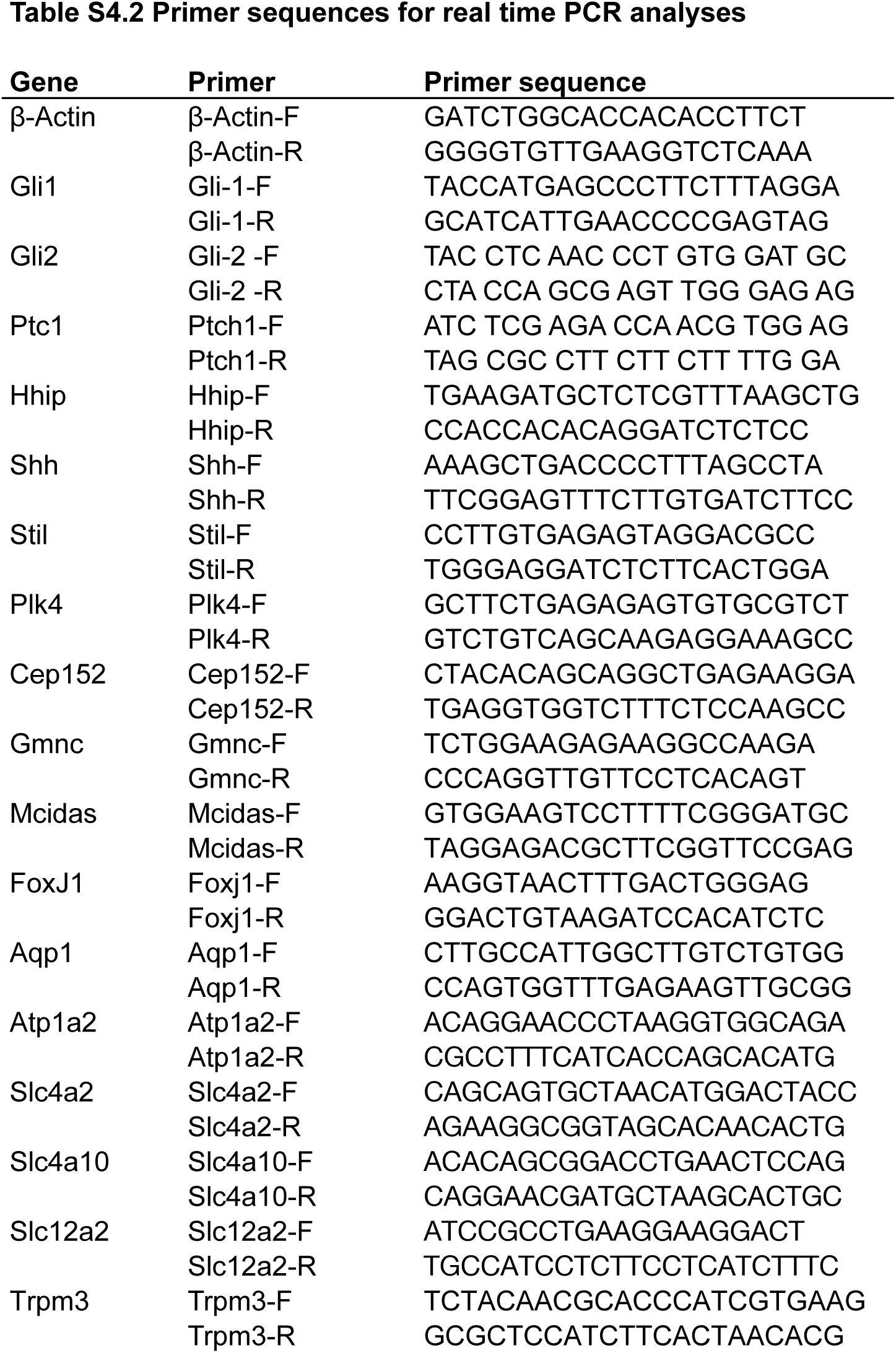
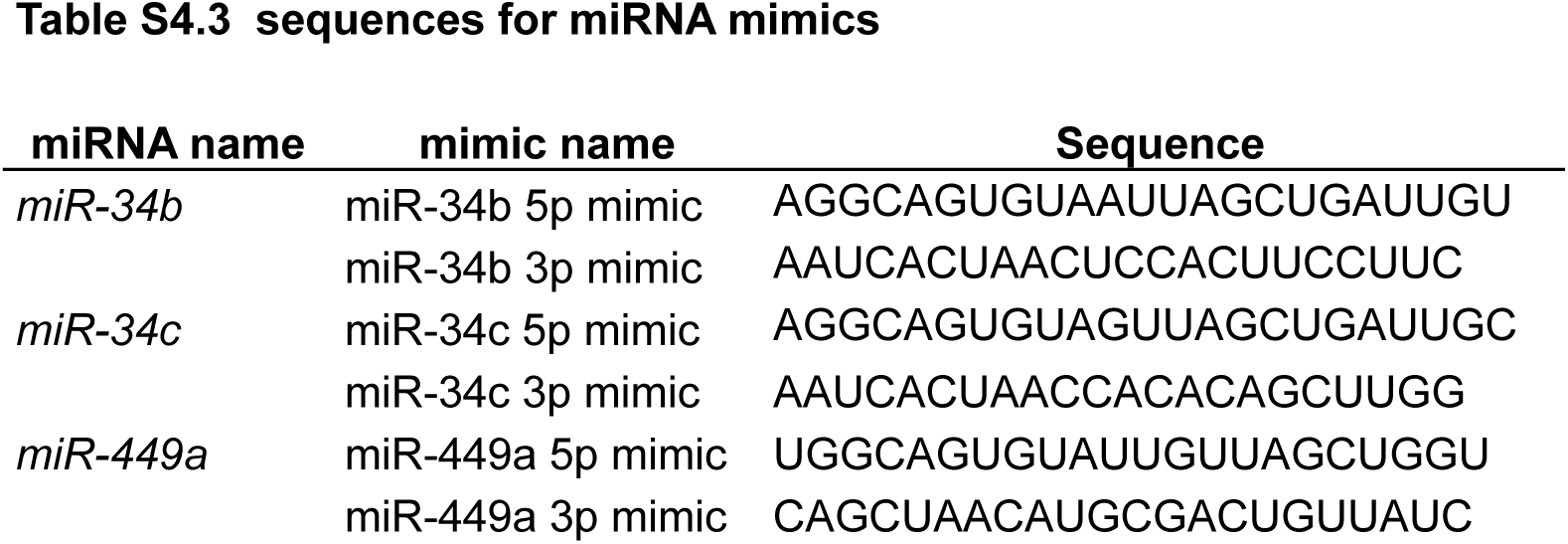
Primer sequences for mouse genotype, qRT-PCR and miRNA mimics.

## Supplementary figure legend

**Supplementary Figure S1.**
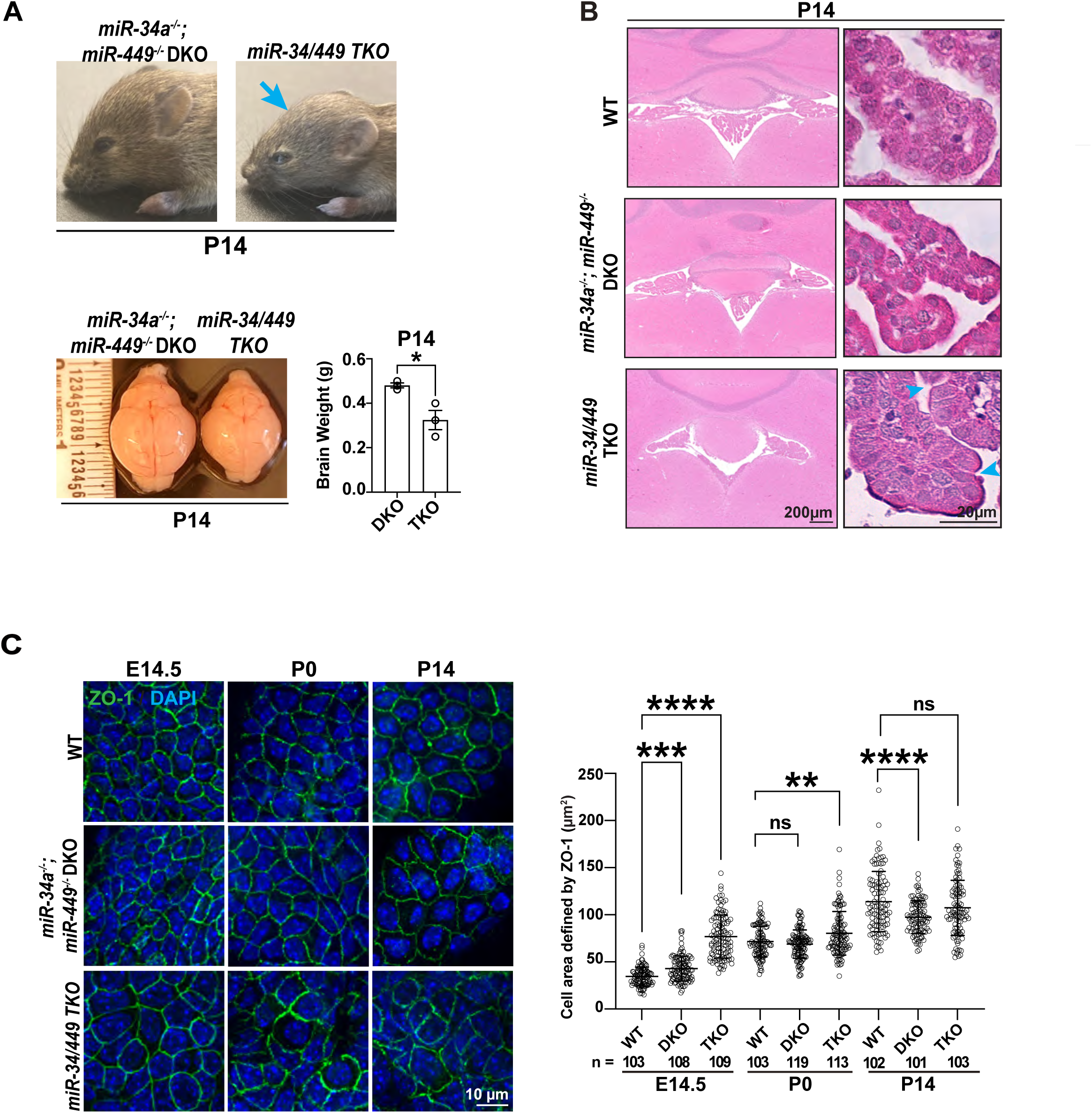
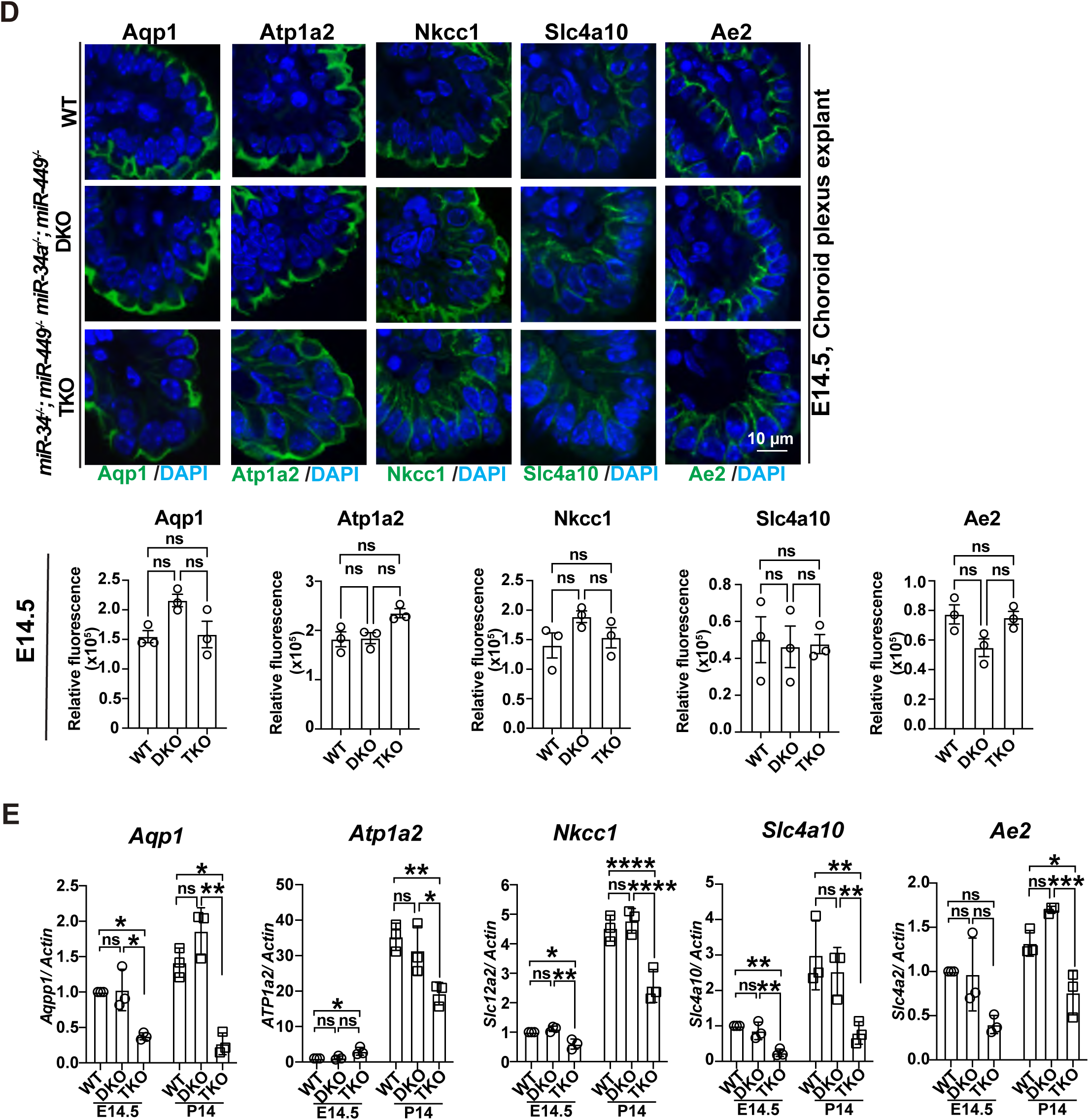
*miR-34/449 miRNAs* deficiency leads to microcephalus and impairs the expression of specific water channels and ion transporters during choroid plexus development. **A.** *miR-34/449* deficiency reduced brain size in postnatal mice. P14 *miR-34/449* TKO mice, when compared with littermate controlled *miR-34a^−/−^; miR-449^−/−^* DKO mice, exhibited a small and narrow forehead, a small brain size, and reduced brain weight. Brain weight quantitation: P14 *miR-34a^−/−^; miR-449^−/−^* DKO vs *miR-34/449* TKO, * *P* = 0.0253, t = 3.483, df = 4; n=3; error bar, sem; unpaired two-tailed Student’s t-test. **B**. *miR-34/449* deficient CPECs exhibited an enlarged cell size with a compact morphology in histology analyses. **C.** CPEC cell size is increased in *miR-34/449* TKO choroid plexus at E14.5 and P0, as quantified by the apical cell area delineated by ZO-1 immunostaining (left). Cell size quantitation: E14.5 WT (n=103) vs. E14.5 *miR-34a^−/−^; miR-449^−/−^* DKO (n=108), \*\*\**P* = 0.0005, t = 3.689, df = 317; E14.5 WT (n=108) vs. E14.5 miR-34/449 TKO (n=109) \*\*\*\**P* < 0.0001, t = 18.78, df = 317; P0 WT (n=103) vs. P0 *miR-34a^−/−^; miR-449^−/−^* DKO (n=119), n.s., *P* = 0.4946, t = 1.062, df = 332; P0 WT (n=103) vs. P0 miR-34/449 TKO (n=113), \*\**P* = 0.001, t = 3.505, df = 332. P14 WT (n=102) vs. P14 *miR-34a^−/−^; miR-449^−/−^* DKO (n=101), **** *P* < 0.0001, t = 4.473, df = 303; P14 WT vs. P14 *miR-34/449* TKO (n=103), n.s., *P* = 0.1102, t = 4.473, df = 303; error bars, sd; one-way ANOVA and Sidak multiple comparisons tests. **D**. *miR-34/449* deficiency does not affect the expression of Aqp1, Atp1a2, Nkcc1, Slc4a10 and Ae2 in E14.5 CPECs. Representative immunostaining images (top) and quantitation (bottom) were shown for Aqp1, Atp1a2, Nkcc1, Slc4A10 and Ae2 for E14.5 wildtype, *miR34a*^−/−^; *miR449*^−/−^ DKO and *miR-34/449* TKO choroid plexus tissue. The statistical tests used in each graph are listed in Supplementary Table S4, n=3; error bars, sd; nested one-way ANOVA and Sidak multiple comparisons tests. **E.** Real time PCR analyses on the gene expression of Aqp1, Atp1a2, Nkcc1, Slc4A10 and Ae2 in E14.5 and P14 WT, *miR34a*^−/−^; *miR449*^−/−^ DKO and *miR-34/449* TKO choroid plexus (n=3 for each genotype). Error bars, sem. The statistical tests used in each graph are listed in Supplementary Table S4, nested one-way ANOVA and Sidak multiple comparisons tests. Error bar, sem.

**Supplementary Figure S2.**
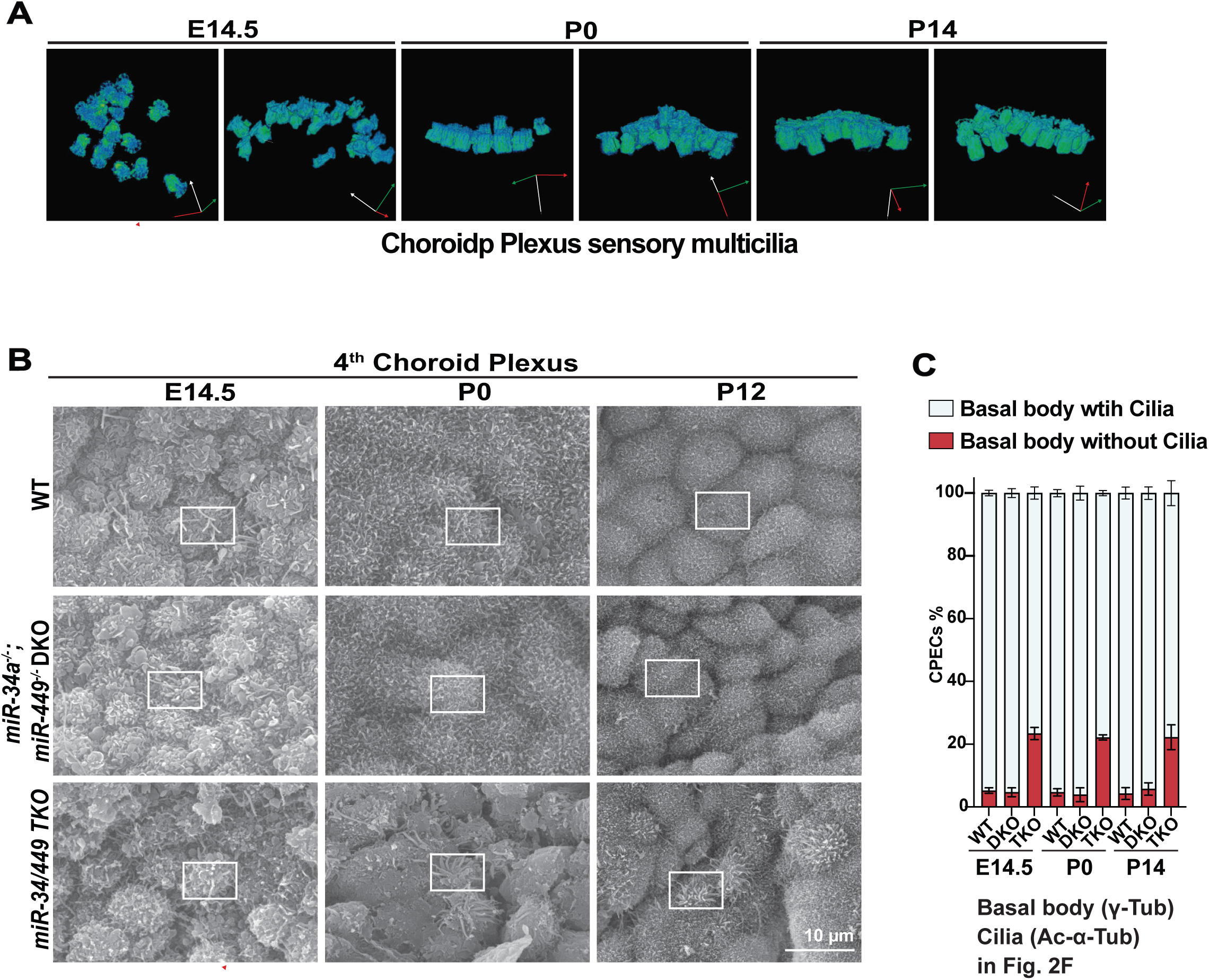
*miR-34/449* deficiency disrupts ciliogenesis in CPECs during embryonic and perinatal development. **A.** Pseudo-color images are shown from FIB-SEM 3D reconstructions of basal bodies in representative E12.5, P0, and P14 WT CPECs. Basal bodies (blue green) are slightly dispersed at E14.5, but are clustered at P0 and become closely aggregated at P14. 3D axes, X (red), Y (green), and Z (white). **B.** SEM analysis of choroid plexus collected from WT, *miR34a*^−/−^; *miR449*^−/−^ DKO and *miR-34/449* TKO mice at E14.5, P0 and P12. White bracelet indicates the area of the zoom-in image in Fig. 2E. **C.** Quantitation of cell number with basal body with or without cilia from immunofluorescence staining in Fig. 2F. in E14.5, P0 and P14 wildtype, DKO and TKO CPECs. Error bar, sd.

**Supplementary Figure S3.**
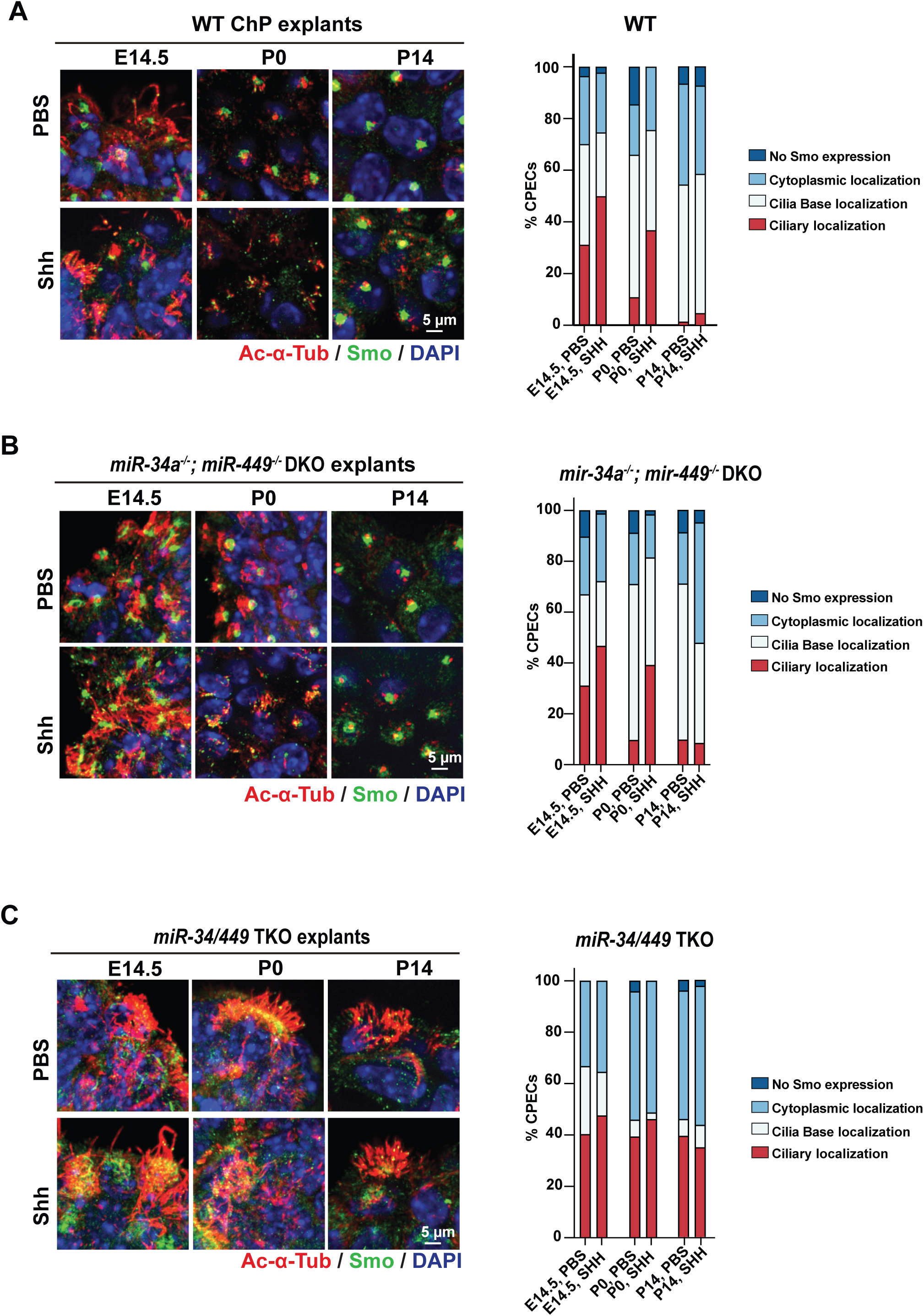
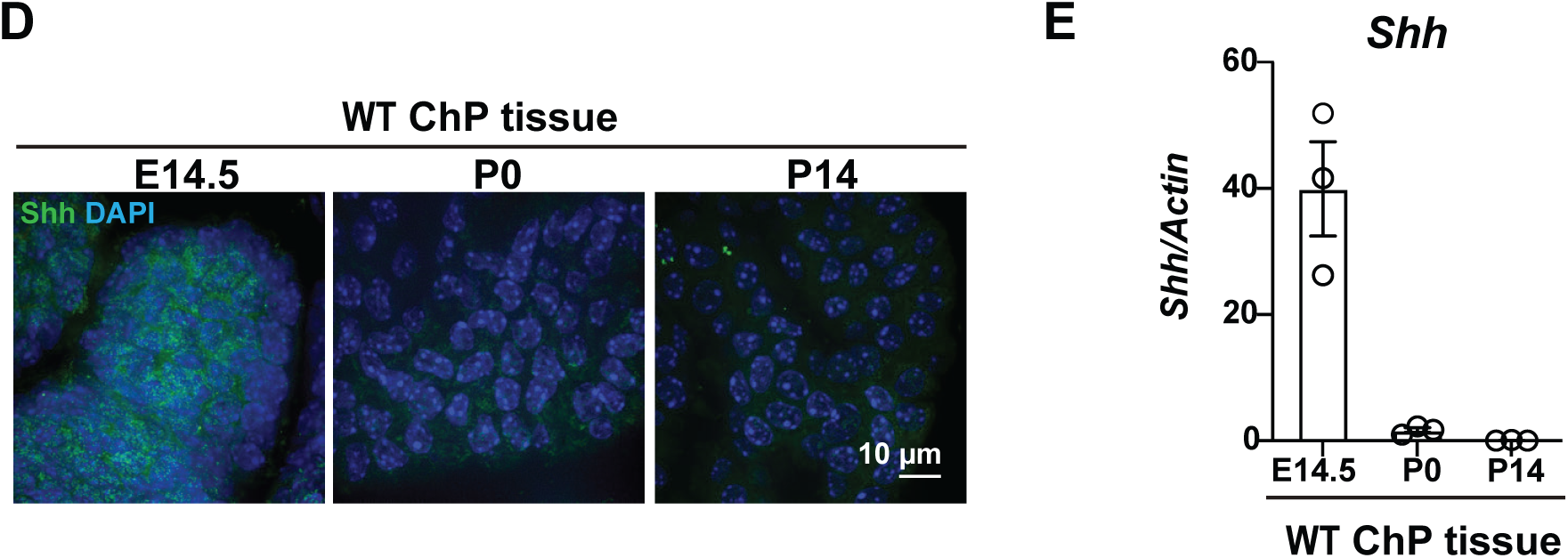
*miR-34/449* deficiency causes aberrant elevation of basal Shh signaling. **A-C.** *miR-34/449* deficiency elevates basal ciliary accumulation of Smo in CPECs. Three pairs of littermate- or age-controlled E14.5, P0 and P14 WT (**A**), *miR34a*^−/−^; *miR449*^−/−^ DKO (**B**), and *miR-34/449* TKO (**C**) choroid plexus explants were treated with PBS or Shh, and subsequently immunostained for Ac-α-Tubulin and Smo. Shh treatment induced ciliary accumulation of Smo in WT and *miR34a*^−/−^; *miR449*^−/−^ DKO CPECs at E14.5 and P0, but not at P14. In contrast, *miR-34/449* TKO CPECs exhibited elevated basal ciliary Smo accumulation at all developmental stages, reaching levels comparable to those of Shh-treated WT CPECs. Exogenous Shh failed to further increase ciliary Smo accumulation in *miR-34/449* TKO CPECs. **A-C**. Representative Smo immunostaining images (left) and quantification of CPECs with ciliary Smo accumulation were shown. **D,E.** Shh protein (**D**) and mRNA (**E**) expression is decreased during wildtype choroid plexus development from E14.5 to P14. n=3 for each developmental stage; error bars, sem.

**Supplementary Figure S4.**
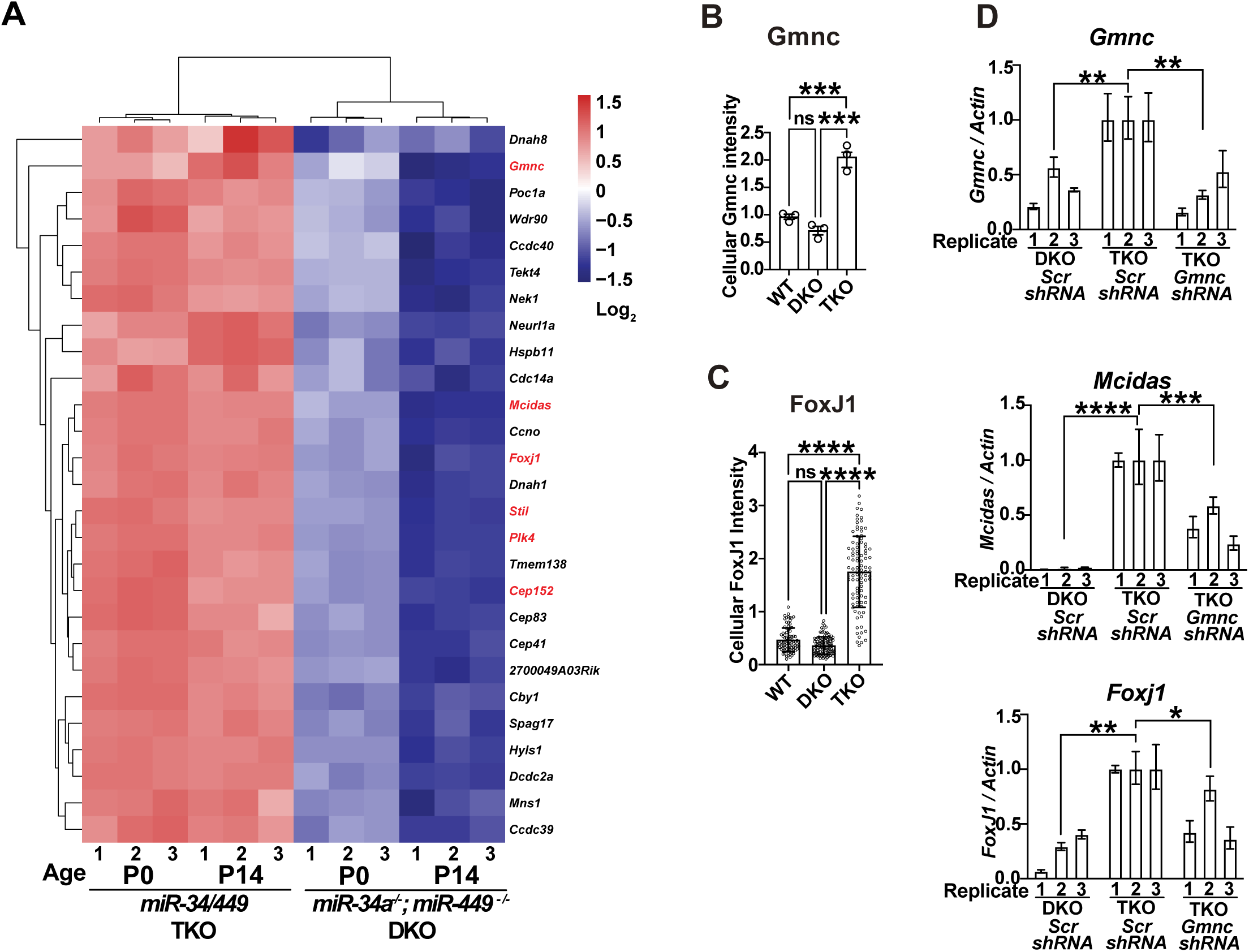
*miR-34/449 miRNAs* regulate FoxJ1-dependent ciliogenesis in CPECs. **A.** A heatmap of RNA-seq data revealed increased expression of motile ciliogenesis genes in P0 and P14 *miR-34/449* TKO choroid plexus compared with littermate-controlled *miR34a*^−/−^; *miR449*^−/−^ DKO samples. Key regulators of multiciliogenesis (Gmnc, Mcidas, and FoxJ1) and basal body assembly (Stil, Plk4, and Cep152) are highlighted in red. **B.** The Gmnc and FoxJ1 immunostaining intensity were quantitated and compared among age-matech or littermate-controlled E14.5 WT, *miR34a*^−/−^; *miR449*^−/−^ DKO and *miR-34/449* TKO mice. n=3 for each genotype. For Gmnc: E14.5 WT vs E14.5 *miR34a*^−/−^; *miR449*^−/−^ DKO, n.s., *P* = 0.294, t = 1.877, df = 6; E14.5 WT vs E14.5 *miR-34/449* TKO, *** *P* = 0.0006, t = 8.105, df = 6; E14.5 *miR34a*^−/−^; *miR449*^−/−^ DKO vs E14.5 *miR-34/449* TKO, *** *P* = 0.0002, t = 9.983, df = 6. For FoxJ1: P0 WT vs. *miR34a*^−/−^; *miR449*^−/−^ DKO, n.s., *P* =0.2298, t = 1.738, df = 280; P0 WT vs. *miR-34/449* TKO, **** *P* < 0.0001, t = 20.56, df = 280; P0 *miR34a*^−/−^; *miR449*^−/−^ DKO s. P0 *miR-34/449* TKO, **** *P* < 0.0001, t = 23.08, df = 280; one-way ANOVA and Sidak multiple comparisons tests. **D.** *miR-34/449* miRNAs target *Gmnc* to reduce FoxJ1 and Multicilin expression in CPECs. P0 *miR34a*^−/−^; *miR449*^−/−^ DKO and *miR-34/449* TKO CPEC monolayer cultures were infected with lentivirus encoding *scramble* (*Scr*) *shRNA* or *Gmnc shRNA*, together with a GFP reporter, and the mRNA level of *Gmnc, Mcidas* and *FoxJ1* were assayed using real time PCR 9 days after infection. *Gmnc* knockdown represses the aberrant increase of *Mcidas* and *FoxJ1* expression in *miR-34/449* TKO CPECs. For the *miR34a*^−/−^; *miR449*^−/−^ DKO *Scr shRN*A vs. *miR-34/449* TKO *Scr shRN*A comparison: *Gmnc*, ** *P* = 0.0043, t = 5.143, df = 6; *Mcidas*, \*\*\*\**P* < 0.0001, t = 12, df = 6; *FoxJ1*, \*\**P* = 0.0038, t = 5.256, df = 6, For *miR-34/449* TKO *Scr shRN*A vs. *miR-34/449* TKO *Gmnc shRN*A comparison: *Gmnc*, ** *P* = 0.003, t = 5.505, df = 6; *Mcidas*, \*\*\**P* = 0.0007, t = 7.302, df = 6; *FoxJ1,* \**P* = 0.0326 t = 3.3, df = 6. n=3 for each genotype; error bars, sd; one-way ANOVA and Sidak multiple comparisons tests.

## Materials and Methods

### Mouse breeding and genotyping

*miR-34a*^−/-;^ *mi-34b/34c*^+/−^; *miR-449*^−/−^ intercross mating were established to generate *miR-34/449* triple knockout (TKO) mice. TKO mice were generated on a mixed genetic background containing C57BL/6, 129 and CD1, and were housed in a non-barrier animal facility at UC-Berkeley (1). All primer sequences for genotype are documented in Supplementary Table S1. All wild-type mice used in this study are C576BL/6J mice (Jax strain #:000664), either purchased commercially or propagated in-house

### Histology analyses

Whole brains were dissected from E14.5, P2 and P14 littermate-controlled wildtype, *mir-34a^−/−^; mir-449^−/−^*DKO, *mir-34/449* TKO, fixed overnight in overnight in 4% PFA (Electron Microscopy Sciences, Cat. # 50-980-495), processed and embedded in paraffin (Fisherbrand^TM^ Histoplast, Cat. #22900700)) by standard procedures. The paraffin sections 5-µm in thickness were stained with hematoxylin (Epredia Signature Series Hematoxylin, Cat. 7211) and eosin (Epredia Richard-Allan Scientific Signature Series Eosin-Y, 7111) to characterize the morphology of brain ventricles and choroid plexus epithelium.

### Real-time PCR analyses

Total RNA was isolated by Trizol (Invitrogen, Cat. #15596) from choroid plexus tissues following the manufacturer’s protocol. For mRNA quantitation, RNA was reversely transcribed using an iScript Advanced cDNA synthesis kit (Bio-Rad, Cat. # 1725038) with random primers. Resulted cDNA was then subjected to SYBR Green-based, real-time PCR analyses (Bio-rad, Cat. # L001894a) using StepOnePlus™a 7900HT real-time PCR system (Applied Biosystems). *β-Actin* was used as an endogenous control. All primer sequences for real-time PCR analyses are documented in Supplementary Table S3. The relative gene expression of genes was plotted by using the (ΔΔct) or 2^−ΔΔct^ method to show the fold change of mRNA expression.

### In situ hybridization (ISH)

Standard histology protocols were used to prepare the P14 whole brain section for *miRNA* ISH using Diethylpyrocarbonate (DEPC) treated water for all procedures (*Song et al*., 2014). After deparaffinization and rehydration, slides were fixed with 4% paraformaldehyde (PFA), treated with Proteinase K, and fixed again with 4% PFA. Slides were incubated first with pre-hybridization solution (3 to 4 hours at 60 °C), and then with hybridization solution mixed with digoxigenin (DIG)-labeled LNA probes against each *miR-34/449 miRNAs* (16 hours at 60 °C). Post hybridization, slides were washed for 10 minutes at 60 °C in a graded series of SSC solutions (2×, 1.5×, 0.2×), then incubated with alkaline phosphatase (AP)-conjugated anti-DIG antibodies in blocking solution. After washing with PBS and alkaline phosphatase (AP) buffer, the slides were incubated with NBT/BCIP in AP buffer to visualize blue ISH signals. Nuclear fast red (Sigma, #N3020) was used for nuclear counterstaining. Slides then were dehydrated and mounted with Permount (Fisher Scientific, #SP15-100). Solutions for ISH (BioChain #K2191020) as described above. DIG-labeled *miR-34a* (ACAACCAGCTAAGACACTGCCA), *miR-34c* (GCAATCAGCTAACTACACTGCCT), and *miR-449c* (CCAGCTAGCAATGCACTGCCT) probes were purchased from Exiqon (#38487-01, #38542-01, and #39641-01 respectively).

### Immunofluorescence staining

For immunostaining on paraffin slides, paraffin-embedded sections, 5um in thickness, were deparaffinized by histoclear (National Diagnostics, Cat. 50-899-90147), rehydrated in a series of ethanol solutions (100% -> 95% -> 90% -> 70%) and water, treated with heat-induced antigen retrieval (Trilogy™, Cell Marque, Cat. 922P-09) and subjected to immunofluorescence staining.

For whole-mount immunostaining, choroid plexus or ependymal tissues were fixed for 2 hours at −20 °C using 80% methanol (EMD, #M×0485P-4) in 20% DMSO (Fisher Scientific, #BP231-100). Fixed tissues were rehydrated in a series of methanol solutions (95%, 90%, 75%, 50%, 25% Methanol) and PBS (PH=7.4), for 5 min at room temperature for each condition, before immunostaining. In both whole mount and paraffin section immunofluorescence staining, rehydrated tissues/slides were blocked in 10% normal goat serum (Vector Laboratories, #NC9270494) in PBS (PH=7.4) for 1 hour at room temperature, incubated at 4°C overnight with primary antibodies (5% normal goat serum in PBS, PH =7.4), washed with PBS for 3 times at room temperature, and incubated with appropriate secondary antibodies (5% normal goat serum in PBS5, PH =7.4) at 4°C overnight. Finally, the samples were counterstained with 4’,6-diamidino-2-phenylindole (DAPI, Vector Laboratories, Cat. #H-1200) at 1µg/mL for 10 minutes at room temperature, before mounting with ProLong^TM^ gold antifade reagent (Invitrogen, Cat# P36930).

Primary antibodies include those against Acetylated-α-tubulin (Sigma, Cat# T6793, 1:300), Arl13b (Abcam, Cat. Ab136648, 1:400), ZO-1 (Invitrogen, Cat. #61-7300, 1:500), Aqp1 (Proteintech, Cat. #20333-1-AP, 1:200), Atp1a2 (Proteintech, Cat. #16836-1-AP, 1:200), Ae2 (Santa cruz, sc-376632, 1:200), Nkcc1 (Proteintech, Cat. #13884-1-AP, 1:200), Slc4a10 (Abcam, Cat. ab122229, 1:200), Smo (Abcam, Cat. ab236465, 1:100), Shh (Proteintech, Cat. #20697-1-AP, 1:200), FoxJ1 (Monoclonal Antibody (2A5), eBioscience™ - 14-9965-82, 1:300), Gmnc (Abcam, Cat. ab86139 1:100). The secondary antibodies include goat-anti-mouse 568 (Invitrogen, Cat# A11031, 1:400), goat-anti-rabbit 568 (Invitrogen, Cat. # A11036, 1:400), goat-anti-rabbit 488 (Invitrogen, Cat. #A11008, 1:400).

### Sample preparation for Expansion microscopy imaging

P0 WT, DKO and TKO choroid plexus explants were subjected to whole-mount immunostaining and prepared for expansion microscopy following the method previously described (2). The monomer solution (4% DMAA (v/v), 34% SA (w/v), 10% AA (w/v), 0.01% Bis (w/v), 1% NaCl (w/v)) in PBS was prepared and stored at 4 °C before use. For gelation, the chemicals 0.001% 4HT, 0.2%APS, 0.25%TEMED and 0.1% methacrolein were added into the monomer solution followed by a gentle vortex. Pre-stained choroid plexus explant were incubated with the gelling solution for 30 min at 4 °C and then transfer to a gelling chamber to incubated overnight in a humidified container at 37 °C for complete gelation.

After gelation, blank gel surrounding the tissue was trimmed and the tissue was then incubated in homogenization buffer (1% w/v SDS, 8 M Urea, 25 mM EDTA, 2× PBS, pH 7.5 at RT) for 6-8h at 80 °C with shaking. Homogenized samples were then washed three times with 1× PBS at RT, followed by at least three washes in 1% decaethylene glycol monododecyl ether (C12E10)/1× PBS. Expanded samples were additionally incubated in 1% decaethylene glycol monododecyl ether (C12E10)/1× PBS at 60 °C for 1 h. Samples were finally washed an additional three times for at least 10 min each with 1× PBS at RT and stored in 1× PBS containing 0.02% sodium azide at 4 °C if necessary. Finally, the gel samples were washed in water for at least 10 min until the sample was fully expanded This was repeated until the sample was fully expanded—at least three exchanges of water before imaging.

### Confocal imaging and quantification

Slides were imaged on spinning disc confocal microscopy (Zeiss LSM 710 AxioObserver or Nikon TE2000-E). All Immunofluorescence images were quantified by using Fiji. Freehand selection was carefully used to select individual cells. Area and integrated density were collected for the measurement. For immunofluorescence on paraffin sections, confocal images were captured as a single slice width at 0.3μm. The integrated density of cellular fluorescence signals of a specific protein, such as Aqp1, was measured by ImageJ for one whole cell, and quantified based on the method previously described (2). Briefly, for each biological sample, 4-6 images were imaged for analysis. The sum of Aqp1/ATp1a2 immunofluorescence intensities for each sample was divided by the number of nuclei (counted by DAPI staining) to quantify the signals for each cell.

For whole mouse immunofluorescence staining, confocal images were Z-stacked for quantification using ImageJ. The apical fluorescence intensity of a protein, such as Aqp1 or Atp1a2, is measured by 7.5 μm-depth (∼25 images along the z-axis, 0.3um per image) stacked images above DAPI-stained nuclei, and DAPI is used as an internal reference across samples as previously reported (*Mao et al*., 2025). The total cellular fluorescence intensity of a protein is measured by a z-stack of all slices stacked in the full depth of a cell and then normalized to that of nuclei (DAPI) as an internal control. Each dot in our quantitation plots represents the ratio of Aqp1/DAPI averaged from 3-6 representative images in a given biological sample. For fluorescence intensity measurement by Image J in each channel, cellular fluorescence intensity = Integrated Density – (Area of selected cell X Mean fluorescence of background readings) (https://theolb.readthedocs.io/en/latest/imaging/measuring-cell-fluorescence-using-imagej.html) For quantitative comparisons, an average of 300∼600 cells were measured for fluorescence intensity in each biological sample/condition.

### MRI-based CSF measurement

Magnetic Resonance Imaging (MRI) was performed using a 7.0 Tesla Bruker Pharmascan system located at the small animal imaging center at UC Berkeley. Neonatal or postnatal mice were anesthetized with isoflurane (Piramal, Cat. 26675-46-7) and positioned in a mouse brain surface array coil (Bruker 2×2) designed for mouse brain imaging. The preparation was inserted into a transmit-only volume excitation coil installed in the magnet. After localizer scans, each mouse was imaged in the coronal and the sagittal views using Rapid Acquisition with Relaxation Enhancement (RARE) pulse sequence, with TR = 5000 ms, TE: T1= = 12 ms, T2 = 60 ms, FOV = 16mm, matrix = 128×128 (resolution = 0.125 mm/pixel), 29 slices = 29 with a thickness of 0.25mm for P7 DKO and TKO mice, RARE factor = 8, bandwidth = 81521.79 Hz. Scan time for each image was 10 min 40 sec. Each image voxel represents 0.0039 mm^3^ in volume. Total CSF volume is measured based on sagittal MRI (µL or mm^3^) = sum of total area (mm2) x thickness (mm).

### Choroid plexus explant culture and treatment

Intact choroid plexus was dissected from the brain fourth ventricles of mice at the indicated developmental stage, washed in PBS, and then transferred onto a permeable membrane (Whatman, Cat# 110614 PC) that floated in the choroid plexus EC media containing DMEM, (Gibco, Cat. #11995-065) with 10% FBS (Hyclone, Cat. #sh30396.03) and 1× Penicillin/Streptomycin (Gibco, Cat. #15140-122). Fresh choroid plexus explants were isolated and then treated with Forskolin (5μM in DMSO, Tocris, Cat. # 1099), recombinant mouse Shh N-Terminus Protein (20nM in PBS, R&D systems, Cat #, 461-SH), or KAAD-cyclopamine (200 nM in DMSO, EMD Millipore Cat. #239804) for 24 hours, before subjecting to experiments as indicated in the figures. Choroid plexus explants treated with PBS or DMSO were used as controls depending on the experiments.

### Choroid plexus *in vitro* primary culture and lentivirus/ AAV infection

P0 DKO and TKO choroid plexus explants were isolated as CPECs at single cell level (3, 4) and cultured on coated 24-well plates. Isolated CPECs were infected with lentivirus stock (titer at 1X10^6^ /mL) with 1:50 dilutions (5, 6) or AAV stock at MOI of 10,000 (7) to 0.4 mL culture media in a 24-well plate for 24h. Infected CPECs were collected 24 h post infection and seeded on permeable, polyester 24-well transwell filter plates coated with collagen for 9-12 days to different experiments and assays when 70% of cells can be transduced by viral infection.

### Luciferase assay

A fragment of the *Gmnc* mRNA 3′UTR containing two predicted *miR-34/449* sites was cloned by XhoI and NotI restriction enzymes into the sites immediately downstream of the stop codon in the Renilla firefly luciferase vector psicheck2 (Promega Cat. C8021). We amplified a fragment of *Gmnc-*3UTR using PCR with *Gmnc*-3′UTR-F, GTTGTTCTCGAGTCCTTACCACACTGCAGATG, and *Gmnc* −3′UTR-R, GTTGTTGCGGCCGCTCAGTAATGCTAAAGGGC. HEK293T cells were cultured in 10% bovine serum in DMEM (Invitrogen, # 11995-073) in a 24-well plate at a density of 5× 10^4^ cells/well. We co-transfected each well of HEK293T cells with 50 ng of psicheck2 constructs, and 100 nM *miR34b, miR34c and miR449a* miRNA mimics (Supplementary Table S1) using *Lipofectamine ™ 3000 transfection reagent* (Invitrogen, #L3000001). At 48 hours after transfection, firefly and *Renilla* luciferase activities were measured using the Dual-Luciferase reporter assay system (Promega, #E1910). The luciferase activity was normalized as the ratio of *Renilla*/Firefly/ luciferase activities.

### Scanning Electron Microscopy (SEM) analysis

Choroid plexus from E12.5, P0 and P14 mice was fixed in Karnovsky’s fixative (Tousimis, Cat. #1011A) in 0.1 M sodium phosphate buffer (Sorenson’s) under room temperature overnight. The fixed tissue was then washed with Sorenson’s sodium phosphate buffer and post-fixed in 1% OsO4 in Sorenson’s for 1 hour. Tissue was dehydrated by passaging through a graded series of ethanol solutions, then critical point dried using a Tousimis 931 Super Critical Point Dryer. The tissue was mounted on aluminum stubs and sputter coated with gold using a PELCO SC-7 coater. The samples were viewed on an FEI XL30 TMP SEM and digital images were collected. SEM was performed in the electron microscopy facility of the University of California at Davis.

### Transmission Electron Microscopy (TEM) analysis

Choroid plexus was harvested from mice and fixed in 2.5% glutaraldehyde/formaldehyde in 0.1M sodium cacodylate buffer, pH 7.2 (EMS, Hatfield, PA, USA) for at least 1 hr. Samples were rinsed (3× 10 min) in 0.1M sodium cacodylate buffer, pH 7.2, (NaCaco) and immersed in 1% osmium tetroxide with 1.6% potassium ferricyanide in 0.1M NaCaco buffer for 1 hour. Samples were then rinsed (3× 10 min) in buffer and then in distilled water (3× 10 min) and then subjected to an ascending ethanol gradient (10min; 35%, 50%, 70%, 80%, 90%, 100%) followed by pure acetone (2× 10 min). Samples were progressively infiltrated while rocking with Epon resin (EMS, Hatfield, PA, USA) and polymerized at 60 °C for 24-48 hours. 70 nm ultrathin sections were cut using a Reichert-Jung Ultracut E ultramicrotome and collected onto formvar-coated 200 mesh copper grids; or 50 nm serial sections were picked up on 1 x 2 mm slot grids covered with a 0.6% Formvar film. Grids were post-stained with 2% uranyl acetate followed by Reynold’s lead citrate, for 5 min each. Sections were imaged using a Tecnai 12 120kV TEM (FEI, Hillsboro, OR, USA) and data was recorded using an UltraScan 1000 or Rio16 cMOS with Digital Micrograph 3 software (Gatan Inc., Pleasanton, CA, USA)(Mcdonald and Webb, 2011).

### FIB-SEM

#### Sample preparation for Focused Ion Beam

Choroid plexus samples were first fixed in 2.5% formaldehyde and 2.5% glutaraldehyde in 0.1M Sodium Cacodylate Buffer, PH=7.4 (Emsdiasum, Cat# 15949), and then stained with a modified osmium-thiocarbohydrazide-osmium (OTO) method in combination with microwave assisted processing, followed by HPF-FS as previously described (8). Briefly, samples were subjected to HPF-FS, and freeze-substituted with 4% osmium tetroxide, 0.1% uranyl acetate, and 5% ddH2O in acetone; then thin embedded and polymerized in Durcupan resin.

Three Durcupan embedded mouse choroid plexus samples (E14.5, P0 and P14) were first individually mounted to the top of a 1 mm copper post, which was in contact with the metal-stained sample for belter charge dissipation (9). A small vertical sample post was trimmed to the Region of Interest (ROI) with a width of 110 µm and a depth of 100 µm in the direction of the ion beam for each sample. The trimming was guided by X-ray tomography data obtained by a Zeiss Versa XRM-510 and optical inspection under a microtome. A thin layer of conductive material of 10- to 20-nm gold followed by 50- to 100-nm carbon was coated on the trimmed samples using a Gatan 681 High-Resolution Ion Beam Coater. The coating parameters were 6 keV, 200 nA on both argon gas plasma sources, 10 rpm sample rotation with 45-degree tilt.

#### FIB-SEM 3D large volume imaging

Three FIB-SEM prepared samples, E12.5, P0 and P14 mouse choroid plexus, were imaged sequentially by a customized Zeiss NVision40 FIB-SEM system previously described (9, 10). Each sample was biased at 400 V to improve image contrast by filtering out secondary electrons. The block face was imaged by a 3 nA electron beam with 1.5 keV landing energy at 1.25 MHz. The x-y pixel resolution was set at 6 or 8 nm. A subsequently applied focused Ga+ beam of 27 nA at 30 keV strafed across the top surface and ablated away 6 or 8 nm of the surface. The newly exposed surface was then imaged again. The ablation – imaging cycle continued about once every two minutes for two weeks to complete FIB-SEM imaging one sample. The sequence of acquired images formed a raw imaged volume, followed by post processing of image registration and alignment using a Scale Invariant Feature Transform (SIFT) based algorithm. The aligned stack consists of a final isotropic volume of 60 x 60 x 60 µm^3^ for each sample, which can be viewed in any arbitrary orientation. Samples E12.5 and P14 were imaged with 8 x 8 x 8 nm^3^ voxel resolution, while sample P0 was acquired at 6 x 6 x 6 nm^3^ voxel resolution throughout entire volumes.

#### FIB-SEM Analysis & Visualization

##### Region of Interest (ROI) selection

All FIB-SEM volumes were first loaded into Fiji, the region of interest (ROI) containing the cilia/basal body region was selected from the tip of cilia to the proximal end of the basal bodies in each cell to generate the 3D volume. Thermo Scientific™ Amira 2019.2-2019.4 (Amira) software was used to navigate the large FIB-SEM data files across the different time domains (E12.5, P0, P14) to identify regions with relevant cellular morphology. The raw data was first visualized in 3D using Amira, and the use of median filters along with partial transparency enabled efficient identification and cropping of ROIs containing relatively high signal-to-noise-ratio basal bodies and cilia. 16 ROIs were selected and cropped in the time domain E12.5. 16 ROIs were selected and cropped in the time domain P0. 17 ROIs were selected and cropped in the time domain P14. Raw FIB-SEM data varied in quality and contrast across ROIs and time domains. 15 ROIs from each time domain were selected for quality and high contrast between the basal bodies and their surroundings. High quality ROIs had the fewest FIB-SEM drift artifacts shearing the cilia or basal bodies and were subjected to further analysis.

##### Segmentation of individual basal bodies and cilia

Each individual basal body was segmented as a secondary ROI. SDA number of each basal body was confirmed by both volume rendering and surface view. Ilastik (version 1.3.3b2) was used to segment basal bodies and cilia from their surroundings(*Berg et al*., 2019). Accuracy and computation time were balanced to produce probability mask files. Using custom MathWorks® MATLAB (R2019a) software scripts the basal body and cilia were separated using their probability mask Ilastik outputs. Results were manually curated for accuracy in Amira, Fiji(Schindelin et al., 2012), and ITK-SNAP(*Yushkevich et al*., 2006). Depending on the severity of missing data in the basal body and cilia probability masks, ITK-SNAP (version 3.6.0) or Ilastik was used for manual image curation.

#### RNA-seq

Using RNA-seq, we characterized dynamic gene expression changes in DKO and TKO in P0 and P14 developmental stages. 100 ng of total RNA from choroid plexus at each stage were used for RNA-seq library construction using Kapa mRNA HyperPrep kit (KAPA Biosysterms, Cat. #, KK8580). The fastq data from RNA-seq were processed by kallisto for quantification(*Bray et al*., 2016) using GENCODE annotation (11). Differentially expressed genes were detected by EdgerR (12). Spearman correlation was calculated to identify genes with consistent increasing or decreasing expression patterns along the developmental time course for choroid plexus (Supplementary Table S1). The heatmap plots produced by pHeatmap (13) in R. Fisher-exact tests were used to measure the significance of the overlapping between gene sets. DAVID bioinformatics tools were used for pathway and function enrichment analysis(14).

#### Quantification and statistical analysis

Statistical analysis was performed using GraphPad Prism (version 10). Appropriate statistical tests were selected based on the distribution of data and sample size. Data are presented as mean ± standard error of the mean (s.e.m) or mean ± standard deviation of the mean (s.d). *P* values in the manuscript present the following star code: ns: p > 0.05 (non-significant),* *P* < 0.05, ** *P* <0.01, *** *P* <0.001, **** *P* < 0.0001. The statistical tests used in each figure are listed in Supplementary Table S4.

